# Temporal single-cell profiling of the parabrachial *Calca* neurons reveals molecular dynamics driving nociplastic pain

**DOI:** 10.64898/2026.08.07.743567

**Authors:** Sekun Park, Hutchinson H. Clarke, Feng Cao, Alexis D. Rose, Eunsoo Yang, Rachel R. Felix, Jonathan Read, Jane Y. Chen, Jordan L. Pauli, Richard D. Palmiter

**Affiliations:** Howard Hughes Medical Institute, University of Washington, Seattle WA 98195 USA; Department of Biochemistry, University of Washington, Seattle WA 98195 USA; Department of Genome Science, University of Washington, Seattle WA 98195 USA

## Abstract

Parabrachial *Calca* neurons are necessary for chronic pain and sufficient to drive nociplastic pain in mice, but how they sustain a pain state that outlasts its trigger is unknown. We performed temporal single-cell mRNA sequencing of the parabrachial nucleus (PBN) across the onset, chronic, and recovery phases of *Calca* neuron-driven tactile allodynia, using fixed-tissue profiling and reference-atlas registration to track molecularly defined populations over time. Activation broadly induced immediate-early genes, after which *Calca* neurons displayed changes in expression of genes that affect signaling and synaptic plasticity, with bidirectional changes that mirrored the onset and resolution of allodynia. The gene encoding brain-derived neurotrophic factor (*Bdnf*) remained persistently elevated in the chronic phase. BDNF infusion in the PBN prolonged allodynia, whereas blockade of its receptor (TrkB) attenuated it, and inactivating the *Bdnf* gene in *Calca* neurons abolished their hyperexcitability, attenuated allodynia, and relieved pain in a migraine model. These observations pinpoint BDNF as a driver of persistent nociplastic pain.

## INTRODUCTION

Tissue damage produces pain that drives protective and coping behaviors. Nerve injury can leave behind chronic, neuropathic pain that outlasts the original insult by months or years^1,2^. Another form of pain, termed nociplastic, can arise without any detectable wound or nerve damage^3–5^. It is rarely localized, tends to spread across the body and is often accompanied by anxiety and depression^3,4,6,7^. Patients also develop allodynia, in which innocuous touch becomes painful (tactile allodynia), and hyperalgesia, in which noxious stimuli feel more severe^3,4^. When nociplastic pain is layered onto another pain condition, conventional analgesics often fail, and treatment becomes difficult^3^.

It is well documented that plasticity in spinal circuits contributes to the transition from acute to chronic pain^1,8–10^. Spinal mechanisms alone, however, cannot account for every case. Tactile allodynia in paw can follow trigeminal nerve damage^11^, and it can be produced by activating neurons in the brain directly, without any peripheral lesion^3,12,13^. Both observations point toward supraspinal mechanisms. Identifying neurons whose activity tracks the pain state would provide a clue for understanding how the brain changes into maladaptive states such as nociplastic pain.

The parabrachial nucleus (PBN) in the dorsal pons is a strong entry point. Anatomical and electrophysiological work identified the PBN as one of the first brain regions to receive nociceptive input from the body^14–19^. PBN neurons project broadly to forebrain targets that mediate the affective and autonomic components of threat responses, including the central amygdala (CeA), the bed nucleus of the stria terminalis (BNST), ventromedial hypothalamus (VMH) and the periaqueductal gray (PAG)^20–24^. They also respond to visceral malaise, bitter taste, hypercapnia, looming cues, and loud sounds^25–31^. This convergence has led to the view that the PBN acts as a general alarm, flagging events that threaten the body regardless of the sensory modality^14,29^.

Within the PBN, several populations have been linked to pain. *Calca*, *Tac1, Nts*, *Tacr1*, *Oprm1*, *Npy1r*, *Pdyn*, and *Slc17a6* neurons respond to transient noxious input^12,21,28,32–35^. Among these, *Calca* (encodes calcitonin gene-related peptide) neurons co-express some of these genes (*Tac1, Nts, Oprm1, Slc17a6*) and they are restricted to the external lateral subdivision. They are necessary for chronic neuropathic pain, and nerve injury enhances their intrinsic excitability^12^. Repeated activation of *Calca* neurons is sufficient to drive long-lasting allodynia without physical injury^12^. Indeed, repeated exposure to most stimuli that activate *Calca* neurons can induce persistent allodynia. Thus, they can generate a diffuse, lasting pain state ─ the defining feature of nociplastic pain ─ making them a tractable model to study.

What remains unknown is how optogenetic or chemogenetic activation of *Calca* neurons leads to a pain state that persists long after the triggering stimulus is gone, which implies lasting molecular changes, but the cells, genes and signals that initiate and maintain synaptic plasticity have not been defined. Here we address these questions with single-cell profiling of the PBN across the onset, chronic, and recovery phases of *Calca* neuron-driven nociplastic pain. The approach yields a time-resolved map of how molecularly defined PBN populations remodel their transcriptomes as a pain state generates and resolves, and it nominates many candidate mediators of persistence. Among them, we focus on brain-derived neurotrophic factor (BDNF) and show that it is both necessary and sufficient to sustain *Calca* neuron-driven nociplastic pain.

## RESULTS

### Temporal single-cell atlas of the parabrachial nucleus across a nociplastic pain states

Persistent allodynia after *Calca* neuron activation outlasts the stimulus by days^12^, so the molecular changes that sustain it should be detectable after the triggering activity dissipates. To capture these changes across the full course of the pain state, we profiled the PBN by single-cell RNA sequencing at three time points. We unilaterally injected AAV-DIO-hM3Dq:mCherry into the right PBN of *Calca^Cre/+^* mice and a few weeks later injected the hM3Dq ligand, CNO, to drive nociplastic allodynia while control animals received AAV-DIO-mCherry. Animals were assigned to an onset group collected 30 min after CNO injection (D0), a chronic group collected 3 days after activation (D3), and a recovery group collected 14 days after activation (D14) (Fig. 1a). At D0, the hM3Dq group showed a significant reduction in paw-withdrawal threshold measured by the von Frey test. This allodynia persisted at D3 and resolved by D14 (Fig. 1b and Extended Data Fig. 1a-c). Chemogenetic activation of the left PBN *Calca* neurons produced a similar time course of persistent allodynia (Extended Data Fig. 1d-f). These data confirm that chemogenetic stimulation of PBN *Calca* neurons induces persistent allodynia that lasts beyond the few hours of CNO-mediated hyperactivity^12^.

**Figure 1.**
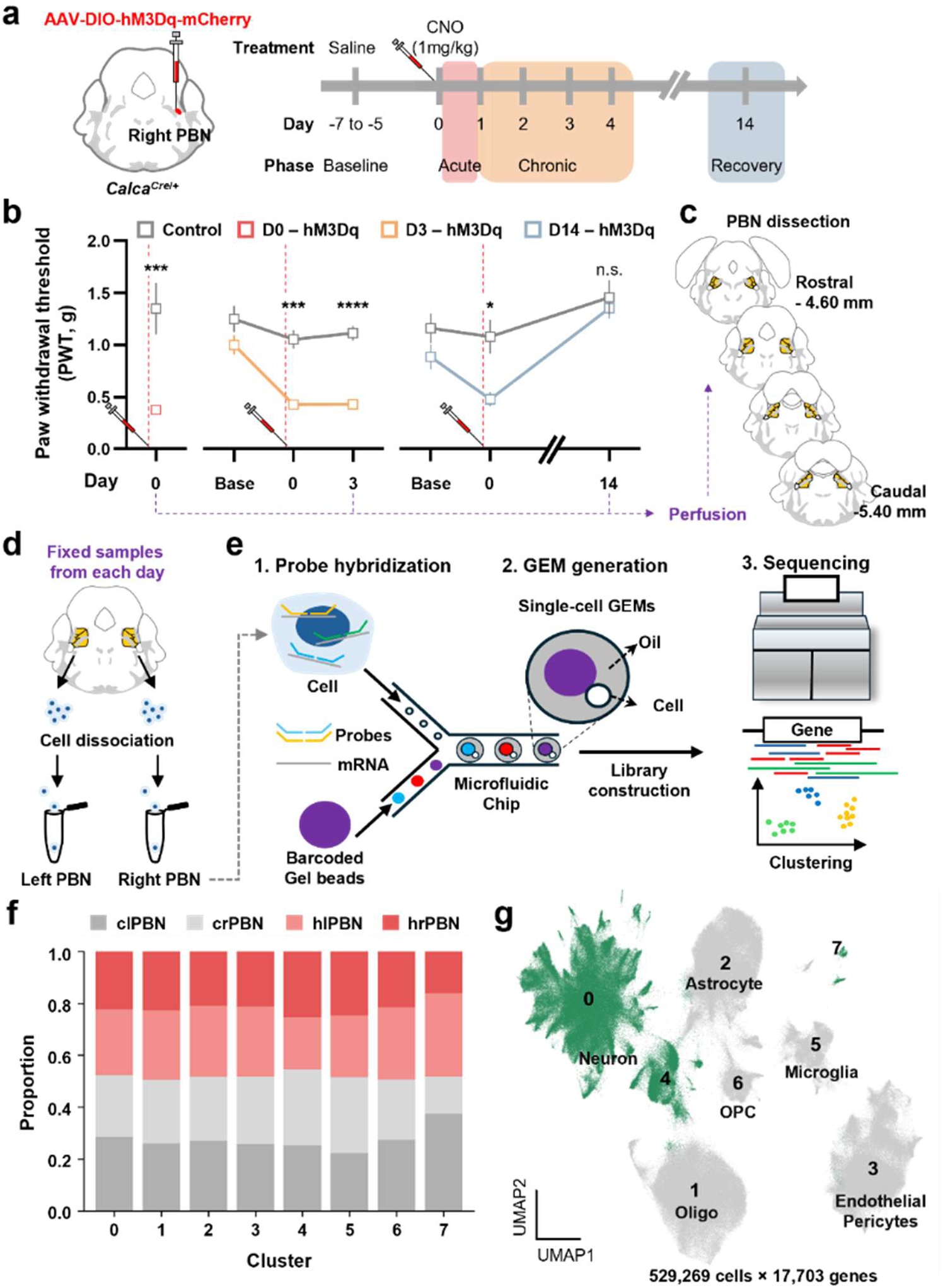
Fixed-tissue, single-cell mRNA sequencing of the PBN across phases of *Calca*-neuron-driven nociplastic allodynia. **a,** Schematic of the *Calca* neuron-driven allodynia model and the time course of the behavioral response. **b,** Paw-withdrawal threshold measured by von Frey filaments at each phase (D0, onset; D3, chronic; D14, recovery). For D0, n = 6 for control and n = 10 for hM3Dq. For D3, n = 8 for control and n = 11 for hM3Dq. For D14, n = 6 for control and n = 7 for hM3Dq. **c,** Rostro-caudal PBN sections included in the analysis. **d,** PBN dissection and cell dissociation. **e,** Single-cell partitioning into gel-beads-in-emulsion (GEMs) and library preparation. **f,** Proportion of each sample across groups and hemispheres within each cluster. **g,** UMAP of all samples showing 8 clusters and their cell types assigned by marker genes. Data are means ±SEM.

At each time point, we collected left and right PBN punches spanning the rostral-caudal axis (bregma -4.60 to -5.40 mm) from experimental and control animals (Fig. 1c and Extended Data Fig. 2a, b). Tissue was perfused, fixed, dissociated, and processed with the Flex gene expression assay (10x Genomics), which halts transcription at fixation and captures mRNAs through hybridized “Mouse whole-transcriptome gene expression” probes (Fig. 1d, e). This fixed-tissue approach let us compare experimental and control cells across all time points within experiments while limiting gene expression artifacts during cell processing. After quality control and doublet removal (Methods), we retained 529,269 cells spanning 17,703 genes across the 12 samples that were evenly distributed (Fig. 1f and Extended Data Fig. 3a). Clustering resolved the major cell types of the PBN region, identified by canonical markers (Extended Data Fig. 3b): neurons (*Thy1*), astrocytes (*Fgfr3*), oligodendrocyte precursor cells (OPCs, *Gpr17*), oligodendrocytes (*Mog*), microglia (*Csf1r*), and a combined endothelial/pericyte population (*Esam*) (Fig. 1g and Extended Data Fig. 3c-j). Sample proportions were similar across clusters, and neurons fell into clusters 0, 4, and 7 (Fig. 1g). This pipeline yielded a quality-controlled, time-resolved dataset of the PBN region.

The PBN punches inevitably include neurons from neighboring regions, so spatial assignment is necessary to restrict the analysis to only PBN neurons. We registered each cell to the Allen Brain Cell Atlas^36,37^ using MapMyCells, which assigns hierarchical Class, Subclass and Supertype labels along with a confidence probability (Fig. 2a). Cells with low mapping Subclass probability (<0.9, 16.0%) were re-classified by supervised clustering (scANVI^38^) in the existing UMAP space (Fig. 2a). The Class distribution matched the initial clustering and included neurons from regions outside the PBN (Fig. 2b). We isolated PBN neurons based on eight PBN subclasses (**217** PB *Lmx1a* Glut; **219** PB-SUT *Tlx3 Lhx2* Glut; **220** PB *Pax5* Glut; **222** PB *Evx2* Glut; **223** B-PB *Nr4a2* Glut; **227** PB-PSV *Phox2b* Glut; **229** PB-NTS *Phox2b* Glut; **265** PB *Sst* Gly-Gaba), resulting in 79,776 cells with 17,703 genes. Most of PBN neurons fell into **217** PB *Lmx1a* Glut and **222** PB *Evx2* Glut, corresponding to the external lateral and dorsolateral PBN regions (Fig. 2c, d). Established molecular markers mapped to the expected subclasses: *Calca* and *Tac1* to **217** PB *Lmx1a* Glut in the external lateral PBN, *Pdyn*, *Foxp2* and *Tacr1* to **222** PB *Evx2* Glut in the dorsolateral PBN, and *Oprm1* broadly across both (Fig. 2e). Independent clustering of the isolated PBN neurons resolved 17 clusters whose marker distribution matched our prior PBN cell atlas^39^, with *Calca* neurons in clusters 1 and 14 (Fig. 2f-g). Atlas registration thus provided spatial identity to the dissociated cells and defined the PBN neuron populations used for all subsequent analyses.

**Figure 2.**
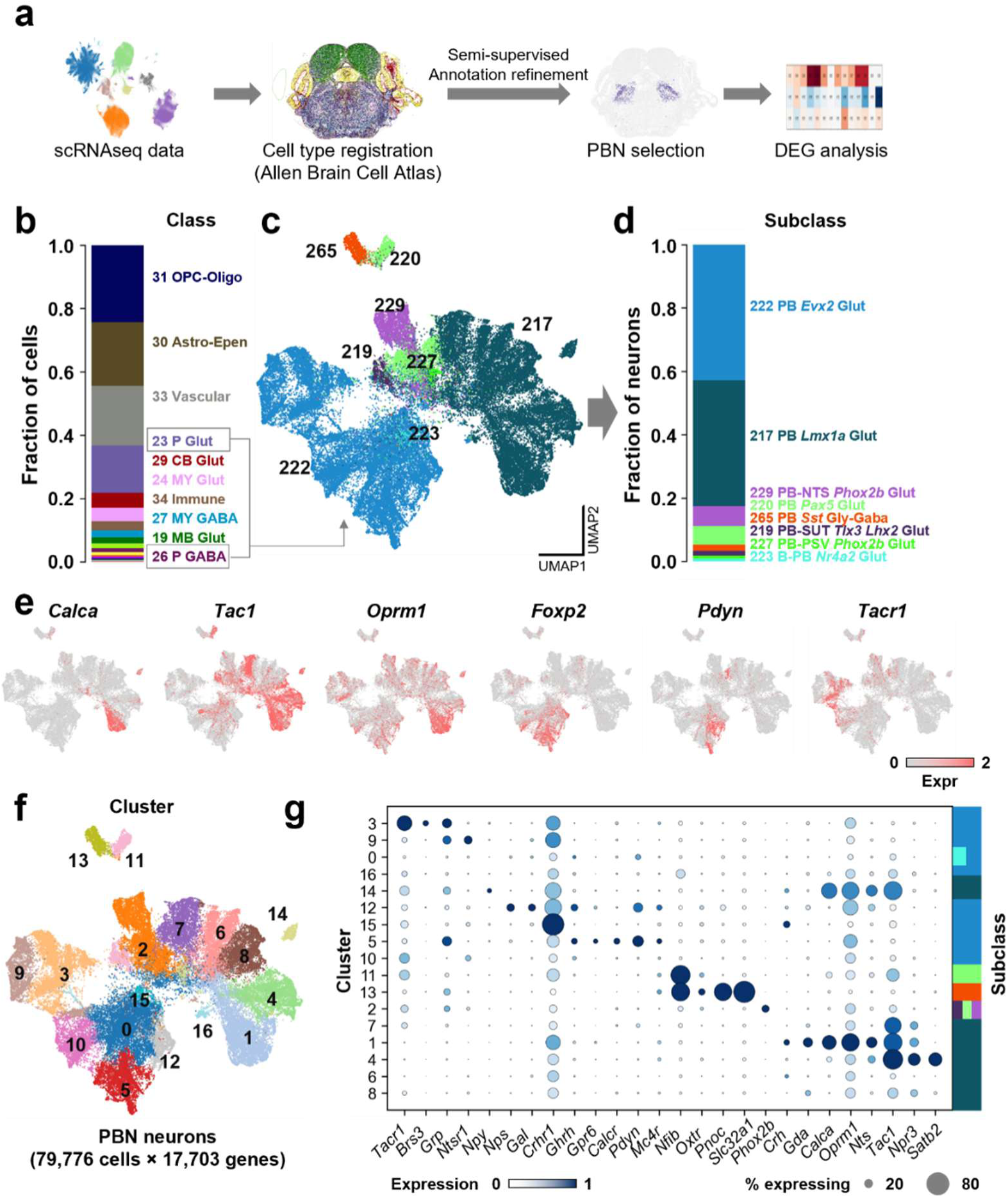
PBN neurons comprise molecularly distinct populations. **a,** Analysis pipeline. **b,** Class distribution registered by Allen Brain Cell Atlas across all samples; PBN neurons were selected for further analysis. **c,** Subclass distribution on the UMAP. **d,** Fraction of each subclass among PBN neurons. **e,** Established PBN marker genes on the UMAP. **f,** UMAP of PBN neuron subclustering. **g,** Marker genes for each cluster, compared with Allen Brain Cell Atlas subclasses (right).

### *Calca*-neuron activation broadly induces immediate-early genes across the lateral PBN

*Calca* neurons were activated at D0 and allodynia persisted to D3, so we asked whether a sustained transcriptional signature accompanied the sustained allodynia, beginning with immediate-early genes (IEGs) as a readout of neuronal activation. We measured the expression of 137 IEGs^40^ as an IEG score, across groups, time points, and clusters in PBN neurons. Then gene expression profiles of **<u>h</u>**M3Dq-expressing **<u>r</u>**ight PBN (hrPBN) were compared to **<u>c</u>**ontrol virus-expressing **l**eft and **<u>r</u>**ight PBN (clPBN + crPBN). On D0, IEG expression rose sharply in the chemogenetically activated PBN (hrPBN) relative to control PBN, confirming that CNO drove neuronal activation (Fig. 3a). This induction was not confined to the clusters containing majority of *Calca* neurons (clusters 1 and 14; Supplementary Table 1). IEG scores also rose in clusters 0-5, 7, and 10-13, indicating that activating *Calca* neurons engages other neurons in the PBN including the GABAergic cluster 13 (Fig. 3b). This increased IEG score disappeared by D3 and D14 (Fig. 3b). To resolve the pattern at single-gene resolution, we examined 12 selected IEGs^41^ that mark neuronal activity, transcription, and synaptic plasticity (*Arc*, *Bdnf*, *Crem*, *Egr1*, *Fos*, *Fosb*, *Fosl1*, *Fosl2*, *Jun*, *Junb*, *Jund, and Myc*). Most were elevated not only in the *Calca* clusters 1 and 14 of the external lateral PBN, but also in clusters 3 and 5, which contain *Foxp2* neurons of the dorsolateral PBN and *Tacr1* neurons of the superior lateral PBN (Fig. 2e and 3c). Thus, CNO-mediated *Calca*-neuron activation promotes broad IEG induction across PBN subdivisions.

**Figure 3.**
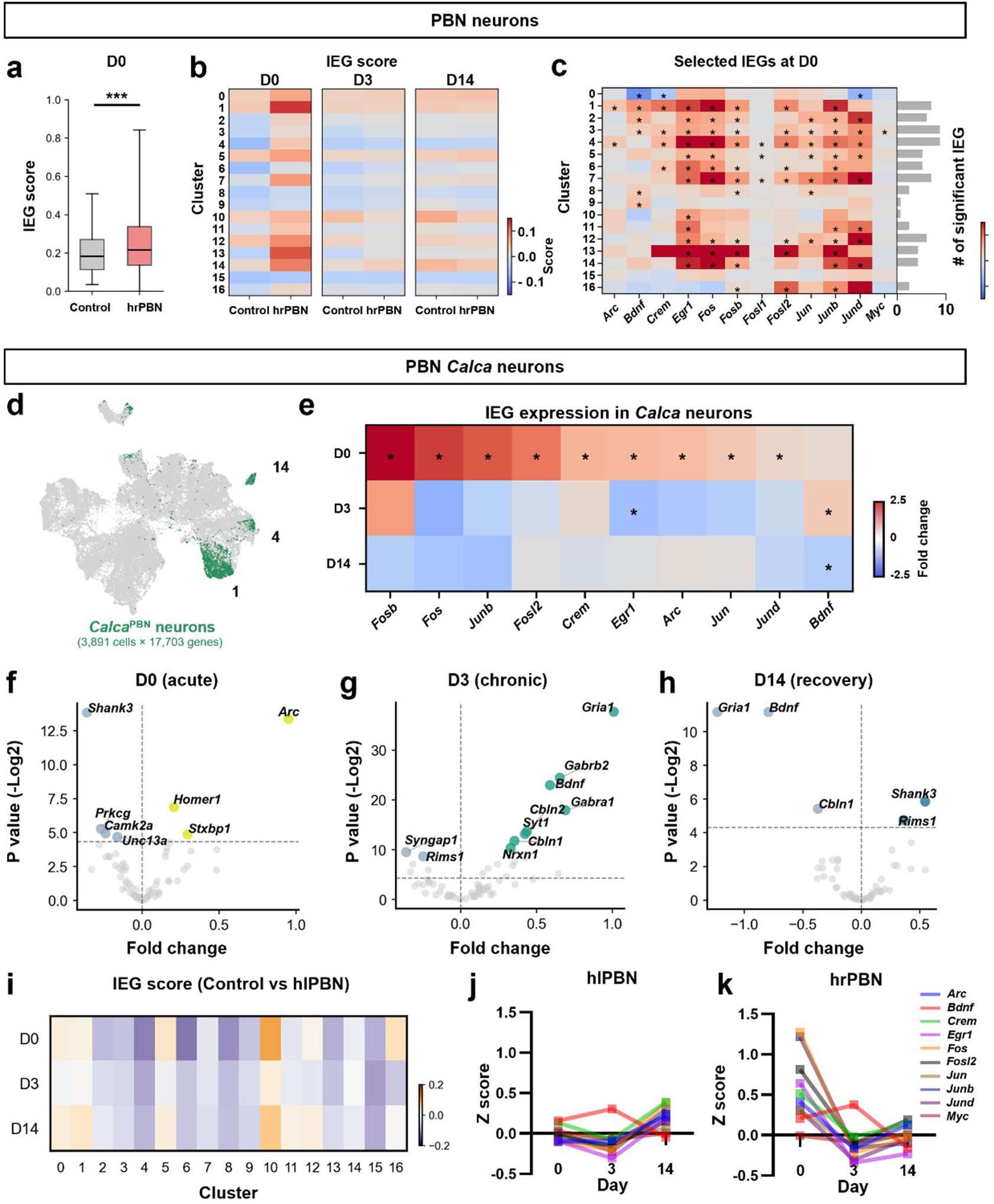
Transcriptional changes in PBN neurons across the time course of nociplastic allodynia. **a,** Immediate-early gene (IEG) score of D0 sample between control PBN (left and right) and the activated hrPBN (right PBN of the hM3Dq group). Black lines in the box plot indicate median and whiskers indicate 95 % CI. **b,** IEG score, control versus hrPBN, across 17 PBN neuron clusters and phases (D0, onset; D3, chronic; D14, recovery). **c,** IEG score distribution among selected IEGs across PBN clusters at D0 (left); bar graph shows the number of clusters with a significant change (right). Control and experimental groups were assessed using a two-sided Mann-Whitney U test. \**P*<0.05. **d,** UMAP of *Calca* neurons (green) among PBN neurons (gray). **e,** Pseudobulk analysis of IEG expression in *Calca* neurons across phases. Control and experimental groups were compared using DESeq2 on pseudobulk profiles (Wald test) with Benjamini-Hochberg-adjusted p values. *Fosl1* and *Myc* are not shown because neither reached significance at any time point. *P<0.05. **f-h,** Volcano plots of differentially expressed genes (DEGs) involved synaptic plasticity and remodeling genes in *Calca* neurons at D0 (**f**), D3 (**g**), and D14 (**h**). **i,** IEG score across phases in control versus hlPBN (left PBN of the hM3Dq group). **j, k,** Z-scored IEG expression across phases in the right (**j**) and left (**k**) PBN of the hM3Dq group.

### *Bdnf* persists while other activity-induced genes return to baseline

The persistence of allodynia past a few hours of CNO-mediated hM3Dq activation^42^ suggested that some may remain active at D3 to maintain allodynia. By pseudobulk differential expression against time-matched controls, *Calca* neurons exhibit 1426, 435, 360 DEGs at D0, D3 and D14, respectively. Among these, most IEGs were elevated 30 min after activation on D0 and returned to baseline by D3 and D14. However, *Bdnf* expression remained elevated through the chronic phase on D3 (Fig. 3e), which nominated it as a candidate for sustaining the pain state after acute activation had subsided.

Because many IEGs are transcription factors, we examined the D0 and D3 data sets for additional genes and programs that could shape synaptic strength and excitability. Among 72 synaptic plasticity and remodeling genes, the D0 response was limited to early-response genes: *Arc* and *Homer1.* The synaptic plasticity-related gene *Stxbp1* was upregulated while *Shank3*, *Prkcg*, *Camk2a*, and *Unc13a* were downregulated (Fig. 3f). By D3, in addition to the sustained rise in *Bdnf*, synaptic plasticity genes *Cbln1*, *Cbln2*, and *Nrxn1* were upregulated along with the AMPA receptor subunit *Gria1* and GABA receptor subunits *Gabra1* and *Gabrb2*, while *Syngap1* and *Rims1* were downregulated (Fig. 3g). The upregulation of *Cbln1* and *Nrxn1* (Fig. 3g and Extended Data Fig. 4a and b) was unexpected because Cbln1/Nrxn1-GluD1 signaling from PBN *Calca*-expressing neurons to the CeA is downregulated in neuropathic pain models^43,44^. The lack of *Ttr* (encoding transthyretin) upregulation in the PBN was also unexpected based enhanced expression in neuropathic pain models^45^ (Extended Data Fig. 4c). We extended this analysis to receptor signaling, examining 78 G-protein-coupled receptor (GPCR)-pathway and 77 receptor tyrosine kinase (RTK)-pathway genes. On D0, *Calca* neurons upregulated the cAMP-associated *Gnas* and *Gnai1* and the receptor *Tacr3*, and downregulated *Sstr1* and *Htr1b* (Extended Data Fig. 5a-c). By D3, GPCRs including *Oprm1*, *Oprl1*, *Gabbr1*, and *Adcyap1r1* were elevated, and within the RTK pathways *Bdnf* again stood out as the gene that remained upregulated at the chronic phase (Extended Data Fig. 5d-f).

For ion channel genes that could affect membrane properties, we analyzed 92 genes across the same conditions (Extended Data Fig. 5g-i). On D0, *Calca* neurons upregulated the potassium channel subunits *Kcnq5* and *Kcnj2* and the chloride channel *Clcn3*, and downregulated *Kcnn2* and *Clcn2*. By D3, more genes were upregulated, including the potassium channel subunits *Kcnj3* and *Kcnk9*, the sodium channel subunits *Scn1b*, *Scn2b*, and *Scn4b*, the calcium channel subunit *Cacng4*, and the acid-sensing channel *Asic1*, while only *Cacna1g* was downregulated. Many of these genes (*Scn1b, Scn2b, Kcna6* and *Kcnk9*) returned toward baseline or were downregulated on D14. The upregulated channel subunits at chronic phase were predominantly those that raise membrane excitability.

A defining feature of these changes was their reversibility. Genes upregulated at D3 returned toward baseline or downregulated at D14. *Bdnf*, *Gria1*, and *Cbln1* were downregulated at recovery, while *Shank3* and *Rims1* rose (Fig. 3h). The molecular state of *Calca* neurons therefore tracked the behavioral course, building during the chronic phase and diminishing after tactile sensitivity returned to normal.

### *Bdnf* induction extends to the contralateral PBN and arises from *Calca* neurons

Although only *Calca* neurons were activated in the right PBN, many IEGs showed activation of non-*Calca* neurons in the right PBN; however, the contralateral *Calca* neurons showed no IEG upregulation in any cluster relative to time-matched controls (Fig. 3i). *Bdnf* was an exception, being significantly upregulated on D3 in the left PBN *Calca* neurons and trending down on D14 (Fig. 3j, k). Broad neuronal activation thus remained confined to the ipsilateral PBN.

The sustained rise in *Bdnf* pointed to *Calca* neurons as its likely source, so we asked whether these neurons express *Bdnf* at baseline. The sequencing data showed *Bdnf* was highest in *Calca* clusters 1 and 14, in unstimulated controls across different days. *Bdnf* mRNA exhibits similar expression patterns with other external lateral markers such as *Nts* and *Tac1* (Extended Data Fig. 6a-c). Using *in situ* hybridization, we confirmed that *Bdnf* mRNA expression was high in the external lateral PBN, and co-expressed *Calca* mRNA (Extended Data Fig. 6d). *Calca* neurons thus exhibit higher baseline of *Bdnf* mRNA in the PBN and it rose further in both hemispheres at D3 even though only right side was activated.

Neuroimmune signaling regulates BDNF expression in the spinal cord^46,47^ and dorsal root ganglia during chronic pain^48^, so we tested whether non-neuronal cells potentially contribute to *Bdnf* in the PBN. We isolated astrocytes and microglia from the dataset and compared activated and control cells in the right PBN. These cells upregulated immune response genes across D0, D3, and D14, including histocompatibility genes (*H2-Q7*, *H2-K1*), and interferon responsive genes (*Bst2*, *Lgals3bp*, and *Zbp1,* Extended Data Fig. 7a-c and e-g). Neither cell type increased *Bdnf* (Extended Data Fig. 7d, h). Non-neuronal PBN cells are therefore unlikely to be the primary source of BDNF after *Calca* neuron activation.

### BDNF signaling in the PBN sustains persistent allodynia

We tested whether BDNF itself maintains allodynia by intracranial infusion into the PBN. We expressed hM3Dq and implanted a cannula over the PBN of *Calca^Cre/+^* mice (Fig. 4a). BDNF infusion (0.5 µg), without *Calca* neuron activation, did not change the paw-withdrawal threshold (Fig. 4b). However, after activation by CNO and establishment of allodynia a day later, BDNF or vehicle was infused into PBN and again 3 days later. The vehicle group recovered by D4, whereas the BDNF group had persistent allodynia (Fig. 4c). We then asked whether BDNF signaling is required for persistent allodynia by blocking the receptor, TrkB with the antagonist ANA-12 (1 µg) infused into the right PBN. ANA-12 given after allodynia was established on D1 had no effect on the ongoing response (Fig. 4e). However, when ANA-12 was infused before *Calca* neuron activation, after CNO administration and again on the following day, persistent allodynia did not develop (Fig. 4f). Hence, BDNF signaling in the PBN by TrkB is necessary for the establishment of *Calca* neuron-driven allodynia.

**Figure 4.**
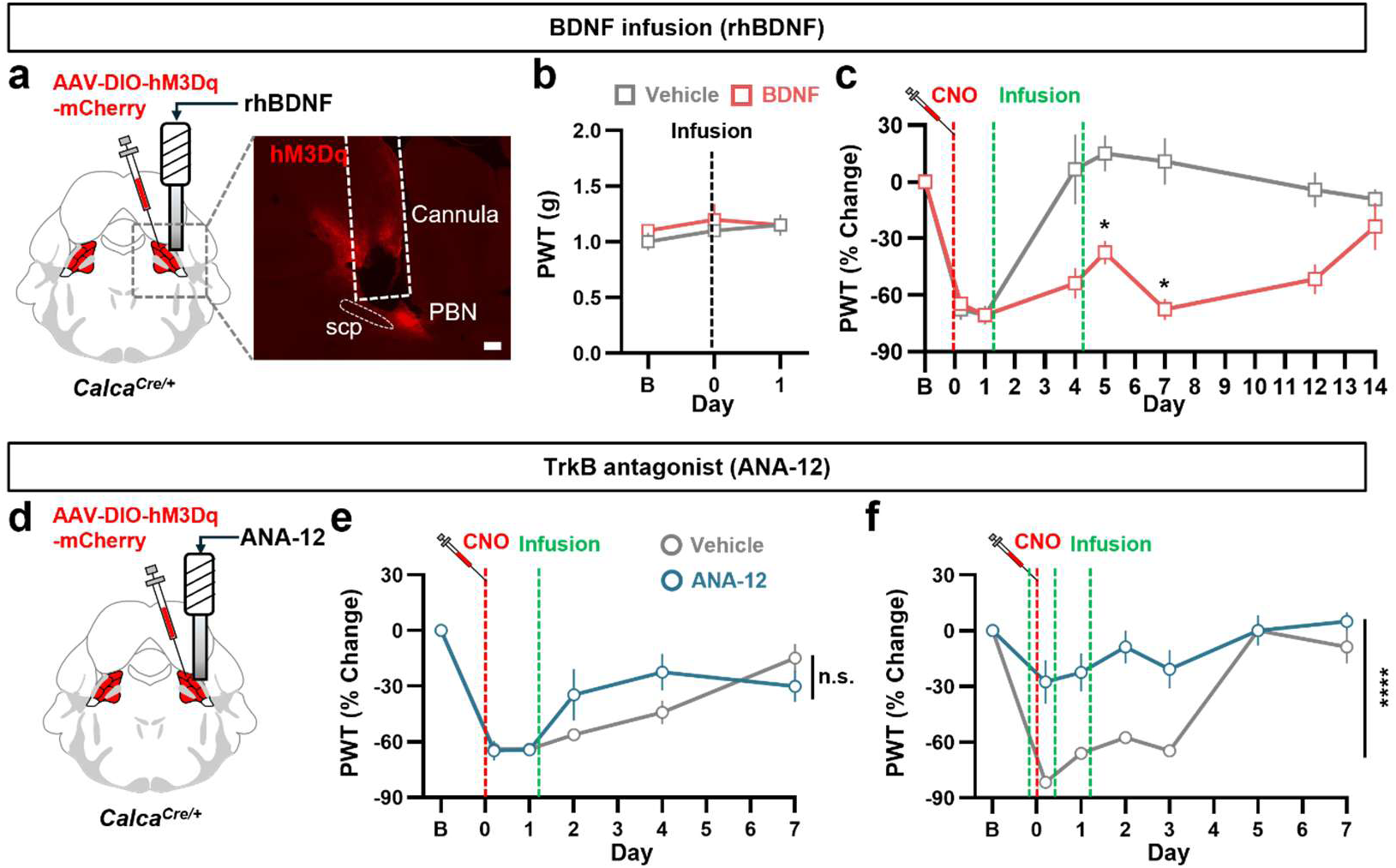
BDNF mediates *Calca* neuron-driven persistent allodynia. **a,** Intracranial BDNF infusion over the PBN (left) and representative image of viral expression of hM3Dq and cannula placement (right). **b,** Paw-withdrawal threshold at baseline (B) and after BDNF infusion. **c,** Paw-withdrawal threshold, as percent change from baseline, across days after *Calca* neuron activation, with BDNF infused on days 1 and 4 after von Frey test (n = 4 for both Vehicle and BDNF group). **d,** Intracranial infusion of the TrkB antagonist ANA-12 over the PBN. **e,** Paw-withdrawal threshold across days after *Calca* neuron activation, with ANA-12 infused on day 1 after von Frey test. **f,** Paw-withdrawal threshold across days after *Calca* neuron activation, with ANA-12 infused three times, one before and two after CNO; n = 4 for both groups. Data are means ±SEM. Two-way Repeated Measured (RM) ANOVA were used to compare groups. Scale bar, 100 µm.

### *Bdnf* from *Calca* neurons drives hyperexcitability and persistent allodynia

To test whether BDNF from *Calca* neurons drives the pain state, we deleted *Bdnf* selectively in these neurons. We crossed a FLP-dependent, *Calca*-restricted Cre line (*Calca^fDIO-Cre/+^*) with *Bdnf^lox/lox^* mice and delivered AAV-FLPo:dsRed bilaterally into the PBN to activate Cre and inactivate *Bdnf* in *Calca* neurons, creating a conditional knockout, cKO (Fig. 5a and Extended Data Fig. 8a-b). We previously showed that chemogenetic activation of *Calca* neurons 3 days in a row increases their intrinsic excitability 2 days after the last CNO by about 2-fold^12^. We used the *Bdnf* cKO mice to test whether it is responsible for the change in intrinsic excitability of *Calca* neurons at D3 (Fig. 5b). In agreement with previous results^12^, *Calca* neurons exhibited regular-firing and late-firing neurons (Fig. 5c). *Bdnf* cKO *Calca* neurons had 38.2% and 48.6% fewer spikes in response to current injection than control neurons in the regular- and late-firing populations, respectively, (Fig. 5d-f), suggesting that *Bdnf* deletion prevents the increase in *Calca* neuron excitability. Input resistance, rheobase, capacitance and resting potential were unchanged (Extended Data Fig. 9a-d).

**Figure 5.**
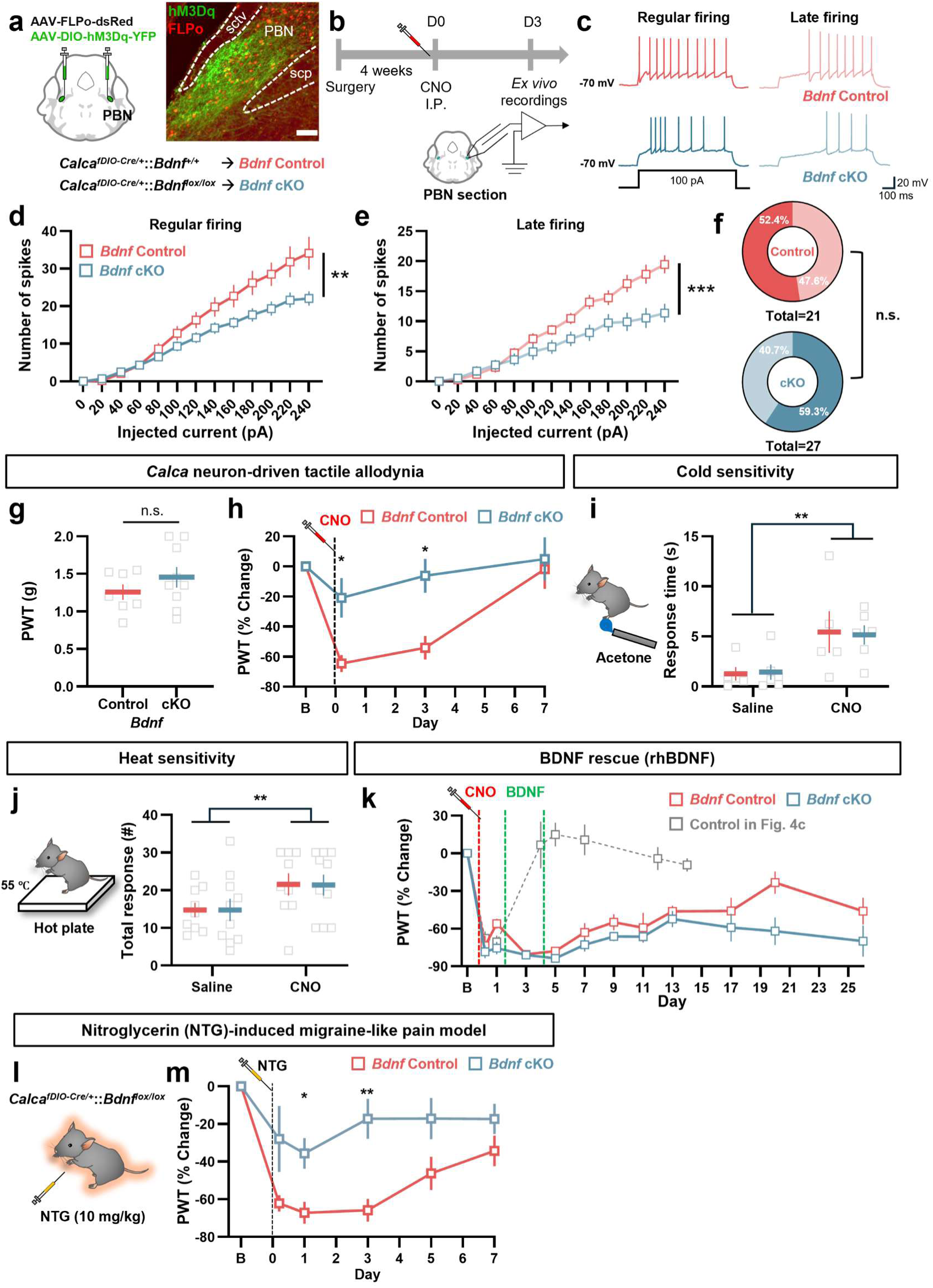
*Bdnf* in *Calca* neurons is required for *Calca* neuron-driven persistent allodynia and NTG-induced migraine-like pain. **a,** Viral strategy for *Bdnf* conditional knockout in *Calca* neurons (left) and representative image of the external lateral PBN (right). **b,** Procedure for whole-cell recording of *Calca* neurons. **c,** Representative action-potential traces of regular and late firing neurons to current injection in *Bdnf* control and cKO *Calca* neurons. **d-e,** Spikes evoked by current injection in regular-firing *Calca* neurons (**d**, n = 10 for *Bdnf* control and n = 16 for *Bdnf* cKO neurons) and late-firing *Calca* neurons (**e,** n = 11 for *Bdnf* control and n = 11 for *Bdnf* cKO neurons) across current steps. Two-way RM ANOVA. **f,** Proportion of regular- and late-firing neurons in *Bdnf* control and cKO groups (n = 21 for control neurons and n = 27 for cKO neurons, Chi-square test). **g,** Basal paw-withdrawal threshold in *Bdnf* control and cKO mice (n = 7 for control and n = 9 for cKO mice, Unpaired t-test). **h,** Paw-withdrawal threshold after *Calca* neuron activation in *Bdnf* control and cKO mice (n = 7 for control and n = 9 for cKO mice, Two-way RM ANOVA). **i-j,** Cold sensitivity by acetone assay (**i,** n = 5 for control and n = 6 for cKO mice) and heat sensitivity by hot-plate assay (**j,** n = 9 for control and n = 10 for cKO mice). Unpaired t-test. **k,** Tactile allodynia after *Calca* neuron activation and the effect of BDNF infusion over PBN in control and cKO mice (n = 4 for all groups, Two-way RM ANOVA.) **l,** NTG-induced migraine model **m,** NTG-induced persistent tactile allodynia (n = 7 for control and n = 10 for cKO mice, Two-way RM ANOVA). Data are means ±SEM. Scale bar, 100 µm.

We then asked whether this loss of hyperexcitability changes pain-related behavior. Conditional deletion of *Bdnf* in *Calca* neurons did not alter basal tactile sensitivity (Fig. 5g). However, after *Calca* neuron activation, the cKO mice displayed reduced allodynia compared to controls (Fig. 5h and Extended Data Fig. 10a-f). Cold and heat sensitivity did not differ between cKO and control mice; and both increased with CNO as expected (Fig. 5i, j and Extended Data Fig. 11a-e). To confirm that the loss of allodynia reflects the absence of BDNF, we tested whether re-supplying it allows development of allodynia. We implanted a cannula over the PBN of cKO mice and infused BDNF on D1 and D3 after *Calca* neuron activation. BDNF infusion produced tactile allodynia that was sustained for 20 days (Fig. 5k). Deleting *Bdnf* in *Calca* neurons therefore attenuates persistent tactile allodynia without affecting baseline or thermal sensitivity.

### *Bdnf* deletion in *Calca* neurons relieves tactile allodynia in a migraine-like pain model

Chronic migraine has features of nociplastic pain, including cutaneous allodynia that spreads beyond the head^7,10,49,50^, so we asked whether *Bdnf* gene expression in *Calca* neurons contributes to this model. We used the nitroglycerin (NTG) model of migraine-like pain, in which *Calca* neurons are activated by NTG and their activity is essential for the pain phenotype^12,51^. We gave NTG (10 mg/kg, i.p.) and measured paw-withdrawal thresholds on the day of injection and on following days (Fig. 5l). The *Bdnf* cKO mice had a much milder and shorter duration decrease in paw withdrawal in response to NTG than control mice (Fig. 5m, Extended Data Fig. 11f and g). This result demonstrates that *Bdnf* expression in *Calca* neurons plays an important role in a pain model that indirectly activates *Calca* neurons.

## DISCUSSION

We performed multiphasic single-cell mRNA sequencing to dissect the molecular changes underlying *Calca* neuron-driven persistent allodynia^12^. Combined with cell-type annotation against the Allen Brain Cell Atlas, this approach resolved the PBN into molecularly distinct populations with spatial resolution^37,39^ and tracked their transcriptomes across the onset, chronic, and recovery phases. Activation of *Calca* neurons induced immediate-early genes both in *Calca* clusters and in other PBN clusters. Among these genes, *Bdnf* stayed elevated through the chronic phase and returned to baseline during recovery. Over the same period, *Calca* neurons changed the expression of genes governing synaptic plasticity and ion channels, and these changes reversed during recovery. We focused on BDNF signaling and found, by infusion over the PBN of BDNF or a TrkB antagonist, that BDNF is sufficient and necessary for persistent allodynia. Selective inactivation of the *Bdnf* gene in *Calca* neurons confirmed that BDNF from these neurons drives both their hyperexcitability and the tactile allodynia.

We implemented a unilateral chemogenetic-stimulation approach because laterality of pain responsiveness has been reported, especially in post-synaptic neurons in the central amygdala (CeA)^23,33,52^. By testing the behavioral response to CNO prior to the sequencing experiments, we could be sure that all the right PBN samples were affected by treatment. We adopted the Fixed RNA Profiling method to capture the earliest changes in gene expression, which revealed robust up-regulation of many transcription factors, primarily of the Fos-Jun family, which presumably orchestrate subsequent events leading to hyperexcitability and prolonged allodynia. While expecting the appearance of IEGs in *Calca* neurons, we were surprised to see that they were also upregulated in many other clusters within the PBN, which could be mediated by neuronal or glial signaling within the PBN or by extrinsic inputs, e.g., from the insular cortex^53,54^.

Analyses of gene-expression changes during onset, establishment, and recovery phases of the allodynia phenotype are a notable feature of the dataset. Genes that were upregulated at the chronic phase returned toward baseline at recovery, and many crossed below it, e.g., *Bdnf*, *Gria1*, and *Cbln1* were downregulated at recovery. Sodium channel and GIRK subunits that rose at D3 fell at D14, while genes such as *Shank3* and *Rims1* moved in the opposite direction. The genes that track this course define a set of candidate mediators of persistent pain that extends beyond *Bdnf.* This pattern suggests that *Calca* neurons may engage homeostatic mechanisms^55–57^ that restore their transcriptional state as allodynia resolves. Our previous study revealed that the duration of tactile allodynia after either chemogenetic or optogenetic stimulation increases from a few days to a few weeks with repeated daily stimulations, but the underlying mechanisms of this scaling phenomenon are unknown^12^.

The upregulation of *Cbln1* and *Nrxn1* in *Calca* neurons and the absence of *Ttr* induction contrast with prior reports for these molecules^43–45^. A likely source of these differences is the distinct means by which chronic pain is induced across models. Those models begin with peripheral nerve injury or inflammation, which alters spinal input and thereby changes the ascending drive onto PBN *Calca* neurons. The *Calca* activation-driven allodynia therefore appears to be established from the top-down rather than from peripheral drive.

We focused on BDNF because BDNF-TrkB signaling is well known to promote synaptic plasticity and could coordinate many of the gene-expression changes we observed^58,59^. Its role in chronic pain has been studied in the dorsal root ganglia (DRG) and spinal cord^48,60–62^, but its role in the brain is less clear and it has not been examined in the PBN^60^. Infusion of BDNF or a TrkB antagonist revealed that BDNF-TrkB signaling is necessary to promote persistent allodynia after *Calca* neuron activation. The data reveal that BDNF synthesis by *Calca* neurons and action in the PBN is important; however, cell-specific inactivation of the *Ntrk2* gene (encodes TrkB) in non-*Calca* neurons will be necessary to establish a paracrine mode of action. BDNF is trafficked by dense-core vesicles and likely released at distal targets where it may also have important functions. From there, it may be retrogradely transported to *Calca*-neuron cell bodies for action^63–65^. TrkB activation engages several intracellular signaling pathways that converge on cAMP-regulated transcription factor, CREB, which may drive the synaptic plasticity that we observed^66,67^.

The hyperexcitability of *Calca* neurons after chemogenetic stimulation may relate to the ion channel changes detected on D3. Upregulation of the sodium channel subunits *Scn2b* and *Scn4b* is consistent with increased sodium current and intrinsic membrane excitability. NaV_β4_ (*Scn4b*) supports resurgent and persistent sodium current, which helps drive repetitive firing^68^. Under acidic conditions, upregulated Asic1 could further depolarize *Calca* neurons through proton-gated currents and add to their excitability, a route separate from the intrinsic changes measured by current injection^69^. How BDNF-TrkB signaling connects to these channel changes is less direct. We detected changes in the auxiliary β subunits (*Scn2b* and *Scn4b*) and BDNF is reported to modulate the α subunit NaV_1.2_ (*Scn2a*)^70^. Because β and α subunits assemble into the same voltage-gated sodium channel complex, the BDNF may enhance sodium current, a plausible route by which BDNF-TrkB signaling could raise the excitability of *Calca* neurons.

By deleting *Bdnf* selectively in *Calca* neurons, we showed that it is required for the persistent allodynia driven by *Calca* neuron activation. This altered sensitivity only occurred when *Calca* neurons were also activated. This dependence indicates that BDNF alone is not sufficient for the underlying plasticity and hyperexcitability, and that *Calca* neuron excitation is also required. This requirement may explain why diverse stimuli that activate *Calca* neurons, including neuropathic pain, visceral malaise, visual/auditory threats, and NTG, all produce persistent allodynia even when the stimulus is unrelated to touch^12^.

In the spinal cord, tactile sensitivity after nerve injury depends on microglia in male but not female rodents^71^. In males, microglia-derived BDNF acts on neuronal TrkB to disinhibit dorsal horn circuits and produce allodynia, whereas females develop comparable allodynia through a microglia-independent mechanism^72^. Our sequencing data pooled both sexes within each sample and therefore do not resolve sex-specific transcriptional responses. At the behavioral level, however, most experiments were not sufficiently powered to detect significant differences (Extended Data Fig. 10c and d) but our prior experiments indicated in minimal sex differences^12^.

We activated *Calca* neurons in the right PBN for most experiments but *Bdnf* rose in the contralateral PBN on D3 without a parallel rise in IEGs. Functional lateralization has been described in the central amygdala across inflammatory, neuropathic, and bladder pain models^23,73,74^, and a recent study extended this to the PBN, where *Tacr1* signaling in the right but not the left PBN is required for neuropathic allodynia^33^. Although these findings point to lateralized pain signaling, *Calca* neuron-driven nociplastic pain itself does not appear to be lateralized: chemogenetic activation of either the right or left *Calca* neurons produced comparable persistent allodynia (Extended Data Fig. 1b), consistent with the bilateral rise in *Bdnf* despite unilateral activation.

Chronic migraine carries features of nociplastic pain, including cutaneous allodynia that spreads beyond the head and comorbid fatigue and sleep disruption. *Bdnf* deletion did not block the acute drop in pain threshold on the day of NTG injection, but it attenuated the persistent phase and facilitated recovery. Peripheral release of CGRP and PACAP is considered to be a major driver of migraine^75,76^, and trigeminal afferents that carry migraine signals project to the PBN via spinal trigeminal nucleus, pars caudalis (Sp5c)^77–79^. We previously showed that NTG activates *Calca* neurons for at least 2 days^12^. Because deleting *Bdnf* in *Calca* neurons relieved the persistent component of NTG-induced allodynia, BDNF acts not only in the *Calca* neuron-driven model but also in the nociplastic component of a disease model.

The CeA is a prominent postsynaptic target of *Calca* neurons, and many of the behavioral and electrophysiological effects of *Calca*-neuron stimulation are mediated by CeA neurons^20,22,23,25,80^. In addition to the direct projection to the CeA, indirect projections to the CeA via the parvicellular ventral posteromedial nucleus, insular cortex and lateral amygdala also display persistent allodynia^81^. Several molecularly defined CeA populations, including *Calcrl*, *Prkcd*, *Sst*, and *Crh* neurons, mediate pain responses by the transient stimulation across chronic pain models^82–85^. Whether these populations contribute to persistent allodynia that we describe or to nociplastic pain such as migraine remains to be determined^3^. Applying the same multiphasic sequencing strategy to the CeA could identify the neural populations that *Calca*-neuron activation engage and the molecular changes within them. The PBN-to-CeA projection likely activates a descending pathway that may involve the periaqueductal grey (PAG) and rostroventral medulla (RVM)^84,86^. Elucidating the details of the circuits and mechanisms by which *Calca*-neuron activation sensitizes pain signaling within the spinal cord remains a challenging endeavor.

## METHODS

### Mouse models

Most experiments used heterozygous *Calca^Cre/+^* mice (JAX 33186) on a C57BL/6 background^87^. *Calca^Flpo^*mice (deposited at JAX) were used for the left PBN activation experiment. *Bdnf* conditional knockout (cKO) mice and wild-type controls were generated by crosses of *Calca^fDIO-Cre^* (see Extended Data Fig. 8) and *Bdnf^lox/+^* mice^25,88^ (graciously provided by Maribel Rios). Animals of both sexes were 8 to 12 weeks old at experimental onset. Mice were group housed with ad libitum access to food and water on a 12-h light/dark cycle at 22°C. After surgery, mice were singly housed. Littermates were randomly assigned to experimental or control groups. All behavioral experiments were performed by people blinded to the experimental details. Viral expression was confirmed histologically at experimental endpoint, and only mice with correct targeting were included in the final analysis. All experimental procedures followed protocols approved by the Institutional Animal Care and Use Committee at the University of Washington.

### Adeno-associated viruses (AAV)

For chemogenetic activation of PBN *Calca* neurons, AAV_DJ_-hSyn-DIO-hM3Dq-mCherry (Addgene) was used for the experimental group and AAV1-Ef1a-DIO-mCherry (in house) for the control group. for chemogenetic activation of left PBN *Calca* neurons, AAV8-hSyn-fDIO-hM3Dq-mCherry (Addgene) was used. For conditional knockout of *Bdnf*, AAV1-CBA-FLPo-dsRed (in house) and AAV1-DIO-hM3Dq-YFP (in house) were co-injected into the PBN. Viral titers were adjusted to approximately 10¹² gc/mL.

### Stereotaxic surgery

Mice were anesthetized with isoflurane and placed on a robotic stereotaxic frame (Neurostar). Virus was injected unilaterally or bilaterally into the PBN (AP: -4.90 mm; ML: ±1.35 mm; DV: 3.40 mm) at 0.1 μl/min for total 0.3-0.4 μl. For intracranial infusion experiments, guide cannulas (RWD Life Science) were implanted above the right PBN (AP: -4.90 mm; ML: ±1.65 mm; DV: 3.2 mm), then secured to the skull with C&B Metabond (Parkell) and dental acrylic. For drug infusion experiments, 26-gauge guide cannulas (RWD Life Science) were implanted unilaterally over the right PBN with same coordinates as above. Mice recovered for 4 to 6 weeks before behavioral testing.

### Fixed-tissue, single-cell mRNA sequencing

Single-cell RNA sequencing was performed using the Chromium Fixed RNA Profiling kit for multiplexed samples (10x Genomics). For preparation of PBN tissue, mice were perfused with 4% formalin (EMS) and 200-µm brain sections were prepared with vibrating microtome (Leica). PBN region was dissected using a sectioning blade (Feather) and tissue punch (Ted Pella). Left and right PBN and CeA were collected in separate tubes. Note that CeA data are collected together with PBN but not presented in this manuscript.

Dissected PBN tissue was fixed overnight, then resuspended in Quenching Buffer (10x Genomics). Cell number and integrity were assessed by propidium iodide staining (Revvity) on a Countess II FL Automated Cell Counter (Thermo Fisher).

For probe hybridization, 50,000 fixed cells per sample were pelleted, combined with a unique Mouse WTA Probe set (10x Genomics). Each biological sample received a distinct Probe Barcode (BC001 to BC004) to enable multiplexing of 4 samples in one reaction tube. Samples were incubated at 42°C for 16 to 24 h in a thermal cycler for hybridization.

After hybridization, cells in each sample were diluted in Post-Hyb Wash Buffer (10X Genomics), counted, and equal numbers from each sample were combined into a single tube. The pool was washed three times to remove non-bound probes, then passed through a 30-μm CellTrics filter (Sysmex). Cell number was determined again before chip loading. Gel Beads-in-emulsion (GEMs) were generated on a Chromium Next GEM Chip Q using the Chromium X instrument (10x Genomics). The pooled sample was combined with GEM Master Mix (10X Genomics), loaded onto the chip. Cells were loaded to target 40,000 cells per GEM well (10,000 per sample). GEMs were incubated in the thermal cycler for probe ligation and barcoding. After GEM recovery, ligated products were pre-amplified by PCR and cleaned with SPRIselect (Beckman Coulter). Sequencing libraries were constructed by sample index PCR, followed by SPRIselect size selection. Library size and concentration were assessed on an Agilent 4200 Tapestation (Agilent Technologies). Libraries were sequenced on an Illumina NovaSeq X by Fred Hutchinson Cancer Research Center Genomics Core.

### Sequencing analysis

Analysis was conducted with a custom Python code. Raw count matrices from each sample were imported into Scanpy^89^ and annotated with metadata for replicate, region, and time point (day). Per-sample quality control retained cells with at least 200 detected genes and removed those flagged as outliers by a 5-MAD (median absolute deviation) threshold on log-transformed total counts, log-transformed gene counts, or the fraction of counts in the top 20 genes. Cells with mitochondrial content above 5% were excluded. Doublets were identified by SOLO^90^ and DoubletDetection^91^, and cells called by both methods were removed. Sample-level QC objects were stored as h5ad files. QC-passed samples were concatenated, and raw counts were preserved in a separate layer. Counts were normalized to 10,000 per cell, log-transformed, and used to identify the top 4,000 highly variable genes with Replicate as the batch key. For initial integration across all cells, PCA was followed by scVI^92,93^ correction on the Sample variable, with neighborhood graph construction, UMAP embedding, and Leiden clustering performed on the scVI-corrected embedding.

Cell-type identification was established using Allen Brain Cell Atlas taxonomy^36,37^. Each cell was assigned class, subclass, supertype, and cluster labels by hierarchical mapping (MapMyCells), and bootstrapping probabilities were stored as confidence metrics. Class and subclass labels were further validated against the Allen taxonomy by using scanVI^38^ to compare neighboring cells’ identification with labeled Allen taxonomy. PBN neurons were extracted based on subclass identity for further analysis. For refined integration of PBN neurons, scVI was trained on raw counts with sample as a categorical covariate and mitochondrial fraction as a continuous covariate. The scVI latent space was used for neighborhood graph construction, UMAP visualization, and Leiden clustering. Cluster-level marker genes were identified by the Wilcoxon rank-sum test (rank_genes_groups) on log-normalized expression, and PBN subclass identity was confirmed using curated markers including Lmx1a, *Calca, Tac1, Tacr1, Oprm1, Pdyn, Foxp2*, and neurotransmitter genes. Differentially expression genes (DEGs) between right and left, control and experimental regions were assessed at each time point (D0, D3, D14) within defined neuronal populations. For immediate-early gene analysis, expression of 137 IEGs^40^ was compared across samples and clusters. From the selected 12 IEGs, pseudobulk analysis was used to identify DEGs between groups. The pseudobulk method was used for following volcano plots for GPCR, RTK and ion channel analysis. Gene lists used in DEG analysis are in Supplementary Table 2.

### Allodynia and hyperalgesia assays

For the von Frey assay, mice were placed in an 11.5 × 7.5-cm chamber with a wire mesh floor and acclimated for an hour for 3 consecutive days before experimental day. In the experimental day, filaments (Bioseb) were applied to the plantar surface of each hind paw 5 times per filament, starting with the 0.16-g filament. Testing ended when two consecutive filaments elicited paw withdrawal in 3 or more of 5 applications. Paw-withdrawal thresholds were averaged across both hind paws because no significant left/right difference was observed across any manipulation. For the hot-plate assay, mice were placed in a 16.5 × 16.5-cm chamber on a 55° C hot plate for 30 s. The total number of nocifensive behaviors (paw flicks, licks, and jumps) was recorded. For the cold assay, mice were placed in an 11.5 × 7.5-cm chamber with a wire-mesh floor and acclimated for an hour. Acetone was delivered by 1.5-inch length tubing connected with syringe containing acetone. A small drop (∼10 µl) of acetone (Fisher Chemical) was applied to each paw and paw flicks, licks and flinching were recorded. Both hot-plate and acetone assay videos were manually analyzed by experimenters unaware of experimental details.

### Locomotion and anxiety assays

For the open field test, mice were placed in a 40 × 40-cm white plexiglass chamber for 30 min. Sessions were recorded with a USB camera, and locomotor activity and center time were analyzed in EthoVision XT18 (Noldus). For the elevated-plus maze test, a plexiglass apparatus consisted of two crossed pairs of arms (two enclosed by 30-cm walls, two open), each 50 cm long and 8 cm wide, elevated 65 cm above the floor. Mice explored the maze for 10 min. Sessions were recorded with a USB camera and analyzed in EthoVision XT 18 (Noldus).

### Pharmacological injections and intracranial infusions

CNO was prepared in sterile saline and administered intraperitoneally: CNO (1 mg/kg; RTI #C929). The recombinant human BDNF (R&D Systems #11166-BD) was reconstituted in sterile PBS. The control group received PBS solution without BDNF. The TrkB antagonist, N-[2-[[(Hexahydro-2-oxo-1H-azepin-3-yl)amino]carbonyl]phenyl]benzo[b]thiophene-2-carboxamide (ANA-12, Tocris #4781), was dissolved in 10% DMSO, 40% PEG300, 5% Tween-80, and 45% sterile saline. The control group received vehicle solution identical to the above but without ANA-12.

For intracranial infusion, BDNF (0.5 µg/µl) or ANA-12 solution were loaded into an injection cannula (RWD Life Science) connected to a microinjector (1 µl total volume). A total of 1 µl of solution was infused at a rate of 8 nl/s (∼ 2 min). Animals were kept connected to the cannula for an additional 5 min in their home cage to allow for diffusion away from the cannula; then the injection cannula was removed and replaced with a dummy cannula.

### Slice electrophysiology

Mice were anesthetized with Euthasol (0.2 mL, i.p.; Virbac) and intracardially perfused with 10 mL of cold (4 to 6°C) cutting solution containing (in mM): 92 N-methyl-D-glucamine, 2.5 KCl, 1.25 NaH_2_PO_4_, 30 NaHCO₃, 20 HEPES, 25 D-glucose, 2 thiourea, 5 Na-ascorbate, 3 Na-pyruvate, 0.5 CaCl_2_, 10 MgSO_4_. Coronal slices (250 μm) were prepared on a vibrating microtome (VT1200S, Leica Biosystems) and recovered for 10 min at 33°C in the recovery solution containing (in mM): 124 NaCl, 2.5 KCl, 1.25 NaH_2_PO_4_, 24 NaHCO_3_, 5 HEPES, 13 D-glucose, 2 CaCl_2_, 2 MgSO_4_ and then transferred to room temperature for at least 45 min. Recordings were performed in ACSF containing (in mM): 126 NaCl, 2.5 KCl, 1.2 NaH_2_PO_4_, 26 NaHCO_3_, 11 D-glucose, 2.4 CaCl_2_, 1.2 MgCl_2_, perfused at ∼2 mL/min at 33°C. All solutions were oxygenated with 95% O_2_ / 5% CO_2_ (pH 7.3 to 7.4, 300 to 310 mOsm). Whole-cell patch-clamp recordings were obtained with a MultiClamp 700B amplifier (Molecular Devices). Only cells with series resistance below 25 MΩ that varied by less than 20% during the recording were included.

For current-injection recordings, patch pipettes were filled with internal solution containing (in mM): 135 K-gluconate, 10 HEPES, 4KCl, 4 Mg-ATP, 0.3 Na-GTP (pH 7.35, 280-300 mOsm). PBN *Calca* neurons were identified by GFP or mCherry epifluorescence and held at -70 mV. Intrinsic excitability was assessed in current-clamp mode by injecting a series of 800-ms current steps ranging from -100pA to 240 pA in 20-pA increments, delivered every 10 s. The number of action potentials evoked at each current step was measured. Data were acquired and analyzed with pClamp 11 and Clampfit 11 (Molecular Devices).

### Fluorescent *in situ* hybridization

Mice were anesthetized with Euthasol (0.2 mL, i.p.; Virbac) and decapitated. Their brains were removed quickly and placed in crushed Dry Ice for rapid freezing and then stored at -80 °C. Coronal sections were cut at 20 μm on a cryostat (Leica), mounted directly onto glass slides (SuperFrost Plus), and stored at -80 °C until staining. RNAscope fluorescent multiplex assay v2 was performed following the manufacturer’s guidelines. Probes used were *Bdnf* (424821-C2) and *Calca* (578771-C3). Five levels of the rostral caudal extent of the PBN were imaged on a Keyence BZ-X710 microscope. Images were captured in a 3 x 3 grid centered on the PBN using the Keyence navigation function and then stacked and stitched in Fiji. Images were minimally adjusted to improve brightness and reduce background.

### Histology and microscopy

Mice were anesthetized with Euthasol (0.2 mL, i.p.; Virbac) and perfused with PBS followed by 4% PFA in PBS. Brains were post-fixed overnight in 4% PFA at 4°C, cryoprotected in 30% sucrose, frozen in OCT compound (ThermoFisher), and stored at -80°C. Coronal sections (30-40 μm) were cut on a cryostat (Leica Microsystems) and collected in cold PBS.

For immunohistochemistry, sections were washed 3 times for 5 min in PBS with 0.2% Triton X-100 (PBST), then blocked for 1 h in 3% normal donkey serum in PBST at room temperature. Sections were incubated overnight at 4°C in PBST containing primary antibodies: chicken anti-GFP (1:10,000, Abcam #ab13970), and/or rabbit anti-dsRed (1:1,000, Takara #632496). Sections were incubated for 1 h in PBS with secondary antibodies: Alexa Fluor 488 donkey anti-chicken and Alexa Fluor 594 donkey anti-rabbit (1:500, Jackson ImmunoResearch). Sections were washed 3 times in PBS, mounted, and coverslipped with Fluoromount-G (Southern Biotech). Fluorescent images were acquired on a Keyence BZ-X710 microscope. Brightness and contrast were minimally adjusted in ImageJ for representation, with identical processing applied across conditions.

### Quantification and statistical analysis

All statistical analyses were performed in Prism 11 (GraphPad). Normality was assessed by the Shapiro-Wilk test to determine whether parametric or non-parametric tests were appropriate. Asterisks denote p values from post hoc tests: *p < 0.05, **p < 0.01, ***p < 0.001. Mice with missed injection sites or sparse expression were excluded from analysis.

## ACKNOWLEDGEMENTS

We thank Susan Phelps and Kit Mandeville for maintaining the mouse colony and lab members for their input during the development of this project. We thank Maribel Rios for sharing the *Bdnf^lox/lox^* line of mice.

## FUNDING

This work was supported by Howard Hughes Medical Institute.

## AUTHOR INFORMATION

R.P. and S.P. designed research and wrote manuscript; S.P., R.F. and J.C. performed fixed-tissue single cell mRNA sequencing. S.P. analyzed sequencing data; S.P., A.R., J.R., H.C., E.Y., performed and analyzed behavioral experiments; F.C. performed electrophysiology experiments; and J.P. performed RNAscope analysis.

## CONFLICTS OF INTEREST

The authors declare no competing interests.

## EXTENDED DATA

**Extended Data Fig. 1.**
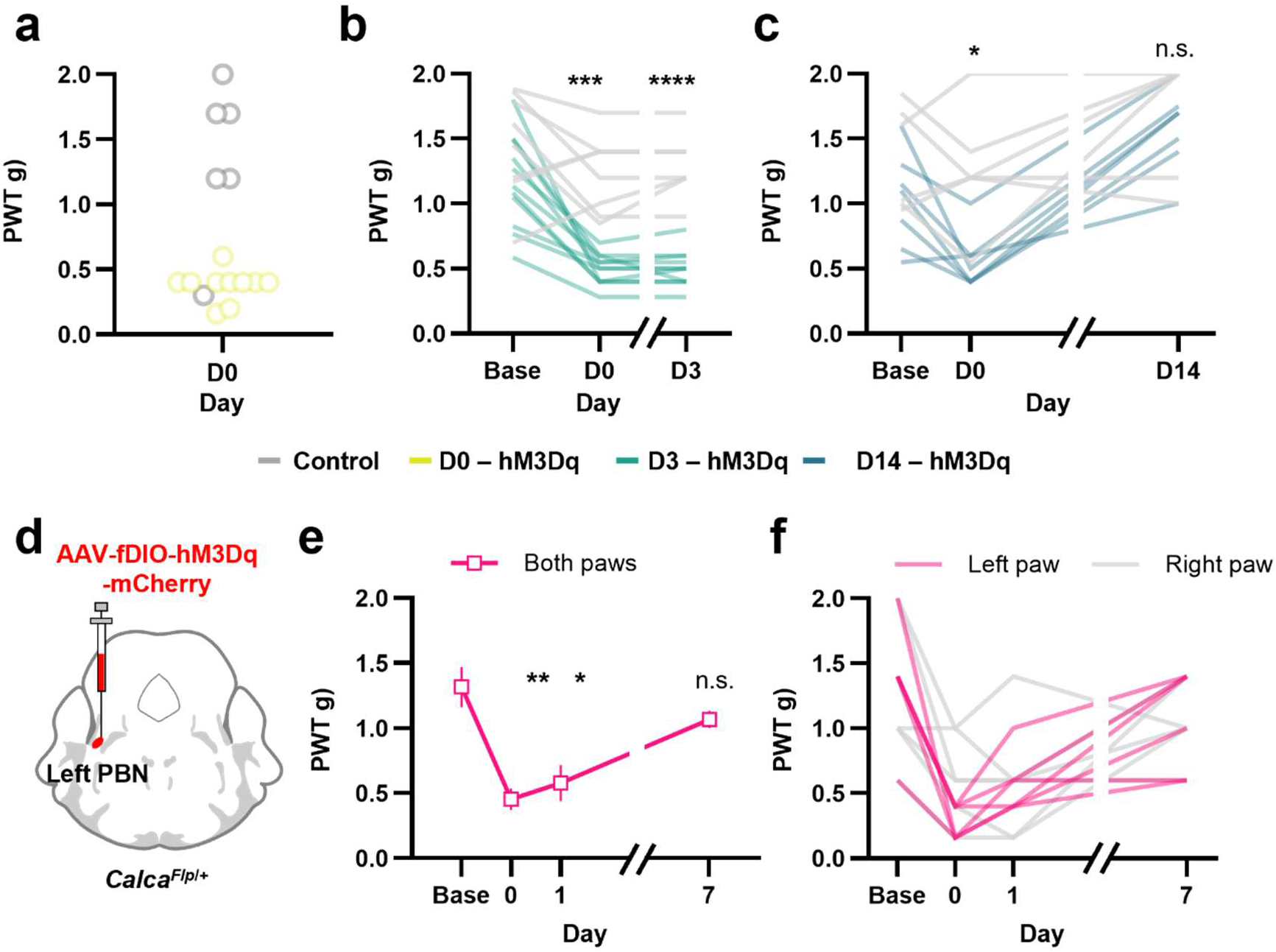
Tactile sensitivity of individual animals in each sample and left PBN stimulation response. **a-c,** Paw-withdrawal threshold (PWT) measured by von Frey test across individual animals of control and hM3Dq group in D0 (a), D3 (b) and D14 (c). For D0, n = 6 for control and n = 10 for hM3Dq (Unpaired t test). For D3, n = 8 for control and n = 11 for hM3Dq (Two-way RM ANOVA). For D14, n = 6 for control and n = 7 for hM3Dq (Two-way RM ANOVA). **d,** Chemogenetic activation of left PBN *Calca* neurons. **e-f,** Paw-withdrawal threshold of both paws (e) and each left and right paw (f); n = 6 for hM3Dq group (Two-way RM ANOVA). Data are means ±SEM.

**Extended Data Fig. 2.**
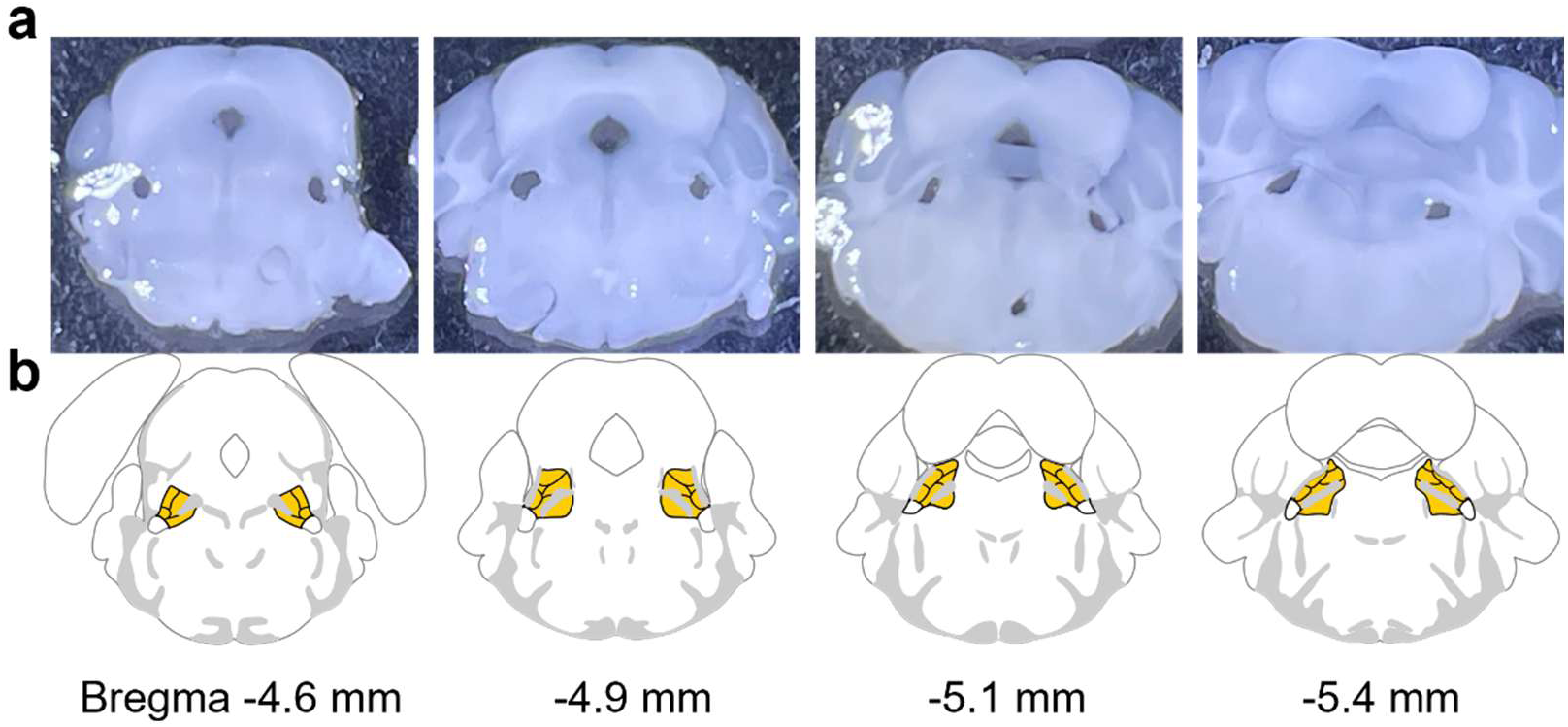
PBN dissection for fixed tissue mRNA sequencing. **a,** PBN sections included sequencing process across rostro-caudal axis. **b**, Corresponding Atlas image and distance from the bregma.

**Extended Data Fig. 3.**
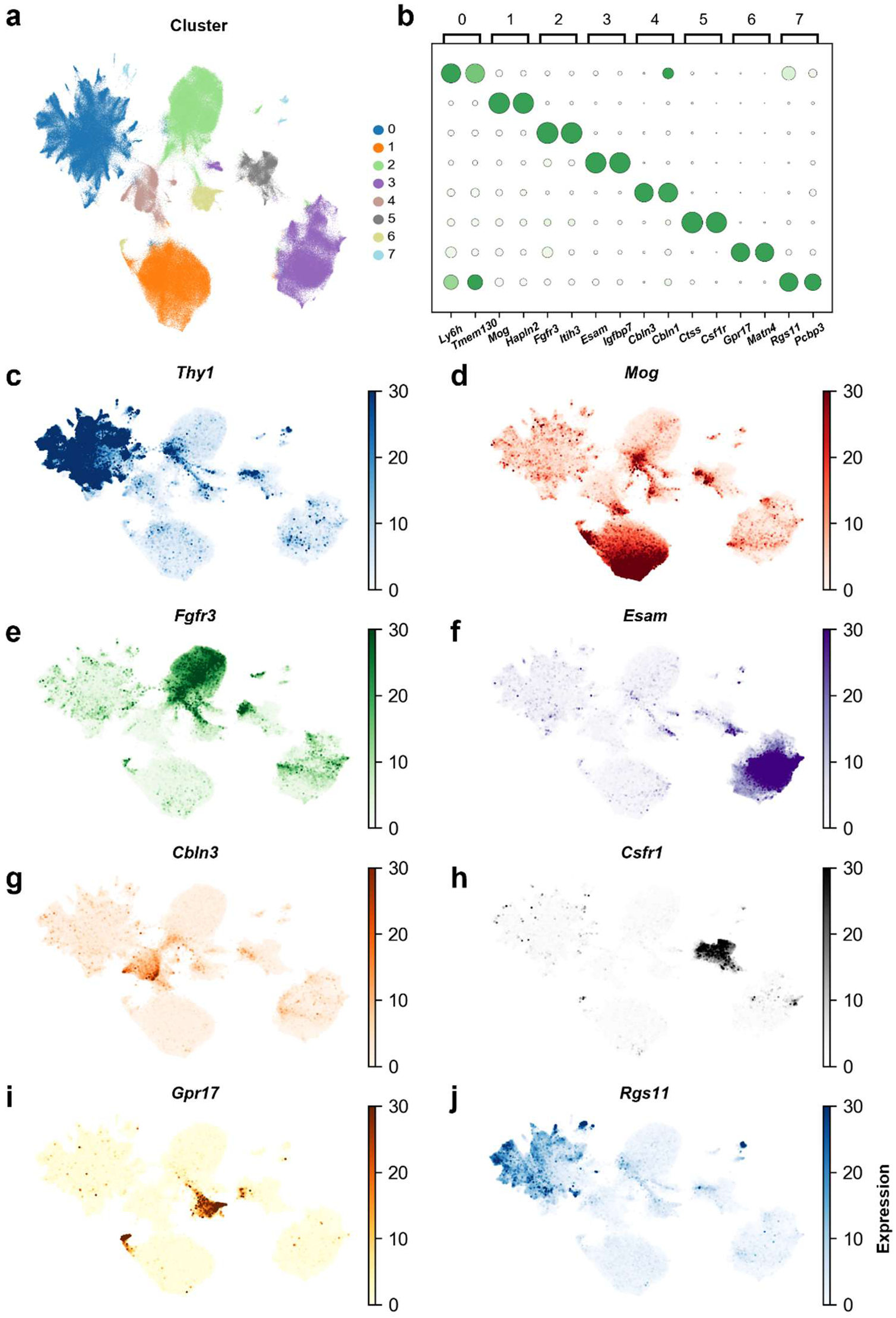
UMAP of distinct population of neuronal and non-neuronal cells across all samples. **a,** UMAP of entire cells with distinct clusters. **b,** Representative marker genes in each cluster. **c-j,** Marker genes expression in UMAP. These marker genes were used to identified neurons (*Thy1, Rsg11*), astrocytes (*Fgfr3*), OPCs (*Gpr17*), microglia (*Csf1r*), oligodendrocytes (*Mog*), and a combined endothelial and pericyte population (*Esam*).

**Extended Data Fig. 4.**
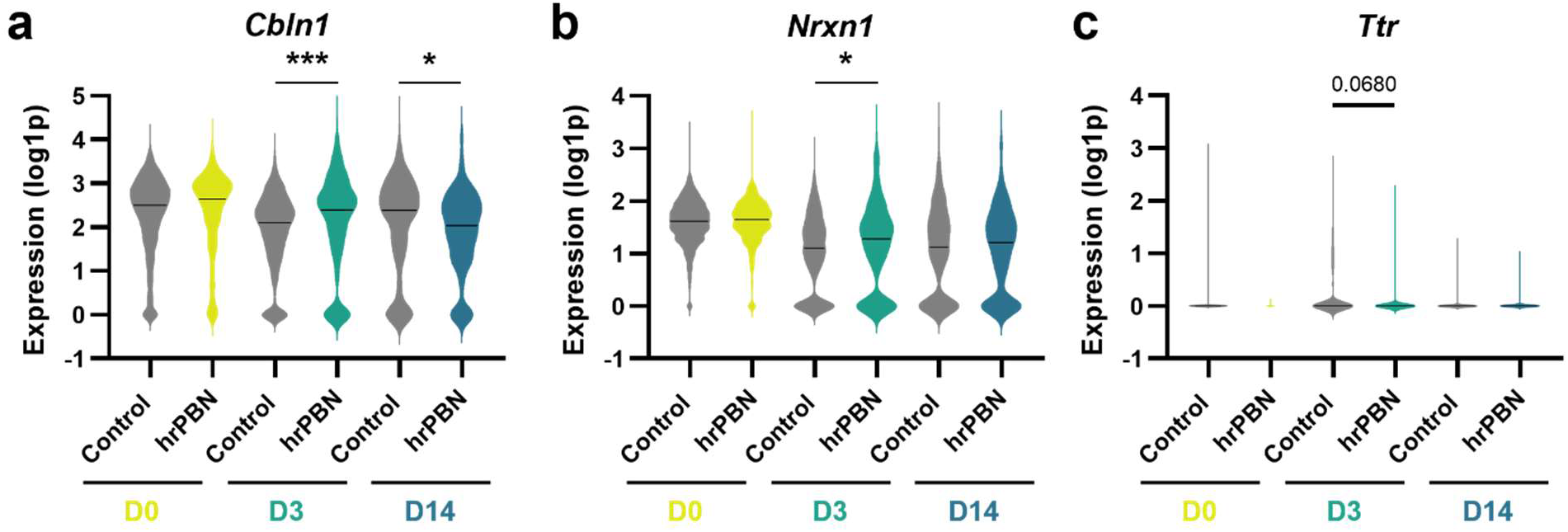
Violin plots of gene expression in PBN *Calca* neurons. a, Cerebellin 1 (*Cbln1*) expression in PBN *Calca* neurons across days. b, Neurexin 1 (*Nrxn1*) expression in PBN *Calca* neurons across days. c, Transthyretin (*Ttr*) expression in PBN *Calca* neurons across days. Black lines in the violin plots indicate the median. Control (clPBN + crPBN) and hrPBN groups were compared by DESeq2 on pseudobulk profiles (Wald test) with Benjamini-Hochberg-adjusted P values. Note that the D0 and D14 comparisons for *Ttr* could not be performed because of the low fraction of *Ttr*-expressing cells. \**P* < 0.05; \*\*\**P* < 0.001.

**Extended Data Fig. 5.**
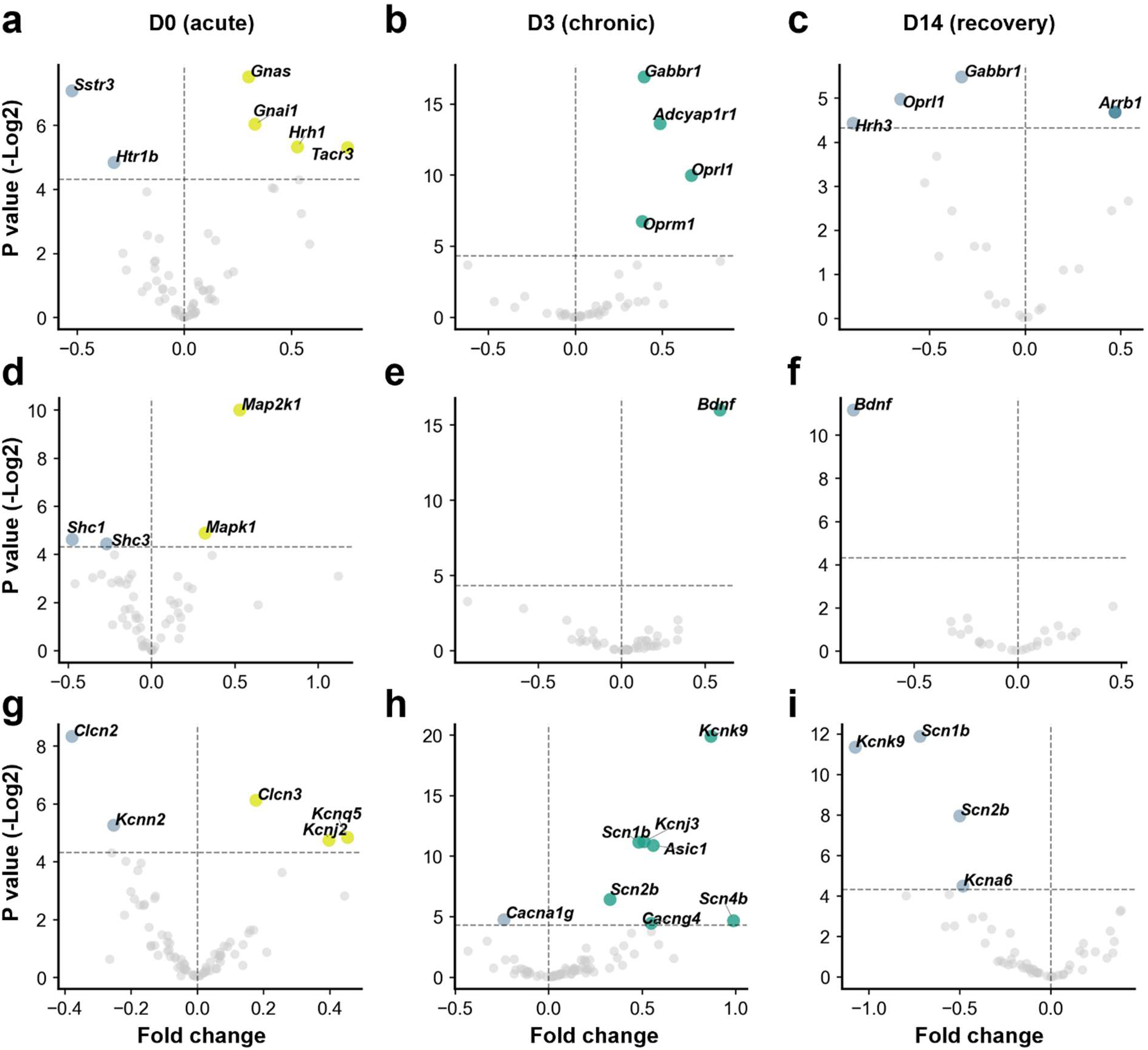
Volcano plots of DEGs in PBN *Calca* neurons across phases of allodynia. **a,** G protein-coupled receptors (GPCRs, 78 genes). **b,** Receptor tyrosine kinase (RTK, 77 genes) pathways. **c,** Ion channels (92 genes).

**Extended Data Fig. 6.**
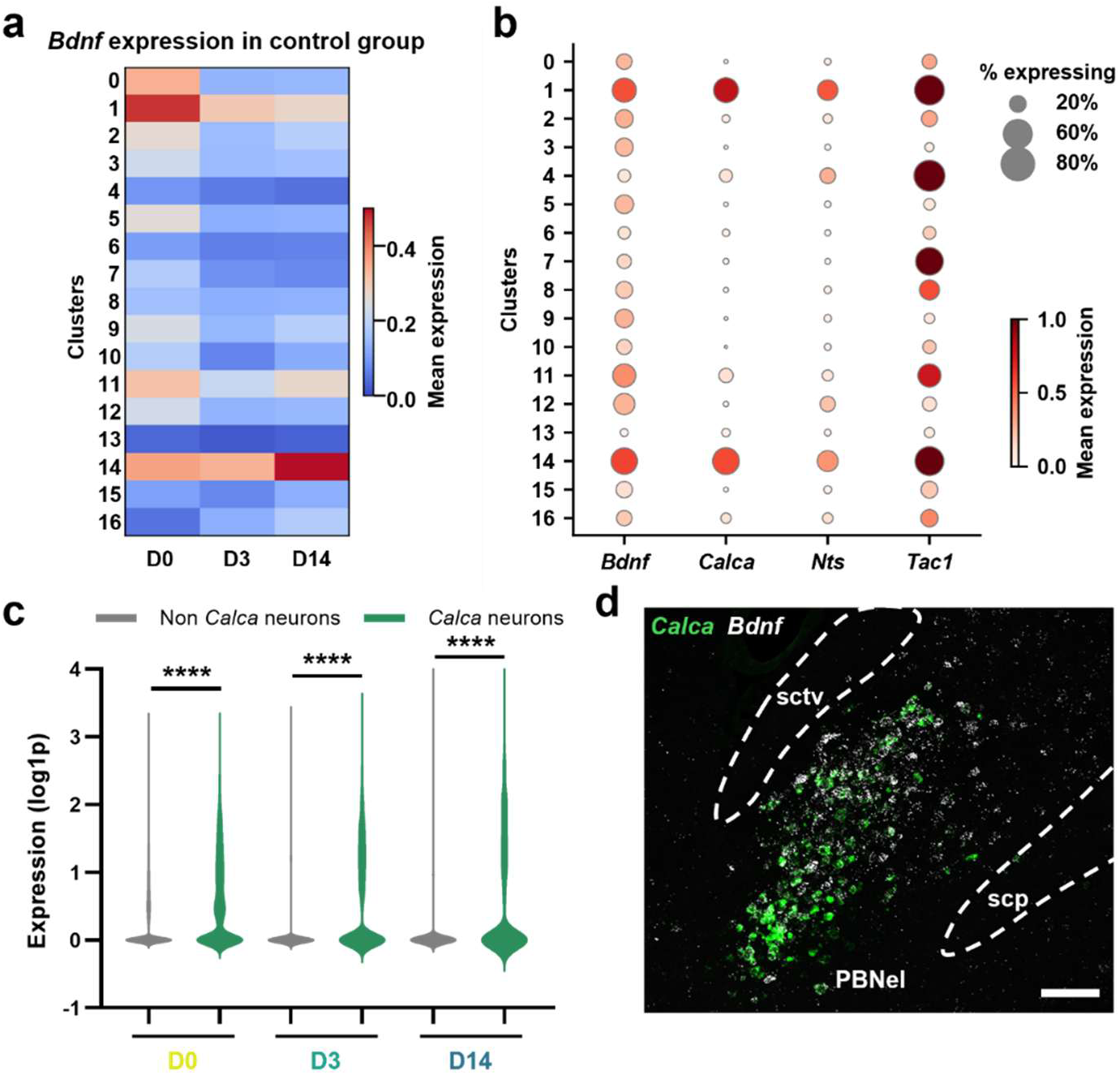
*Bdnf* expression in the PBN of the control group. **a,** Heatmap of *Bdnf* expression in the control group across days and clusters. **b,** Dot plot comparing *Bdnf* expression with other external PBN markers. **c,** Violin plots of *Bdnf* expression in *Calca* versus non-*Calca* neurons. Groups were compared by DESeq2 on pseudobulk profiles (Wald test) with Benjamini-Hochberg-adjusted P values. **d,** Fluorescence *in situ* hybridization image of *Bdnf* and *Calca* mRNA in the external lateral PBN.

**Extended Data Fig. 7.**
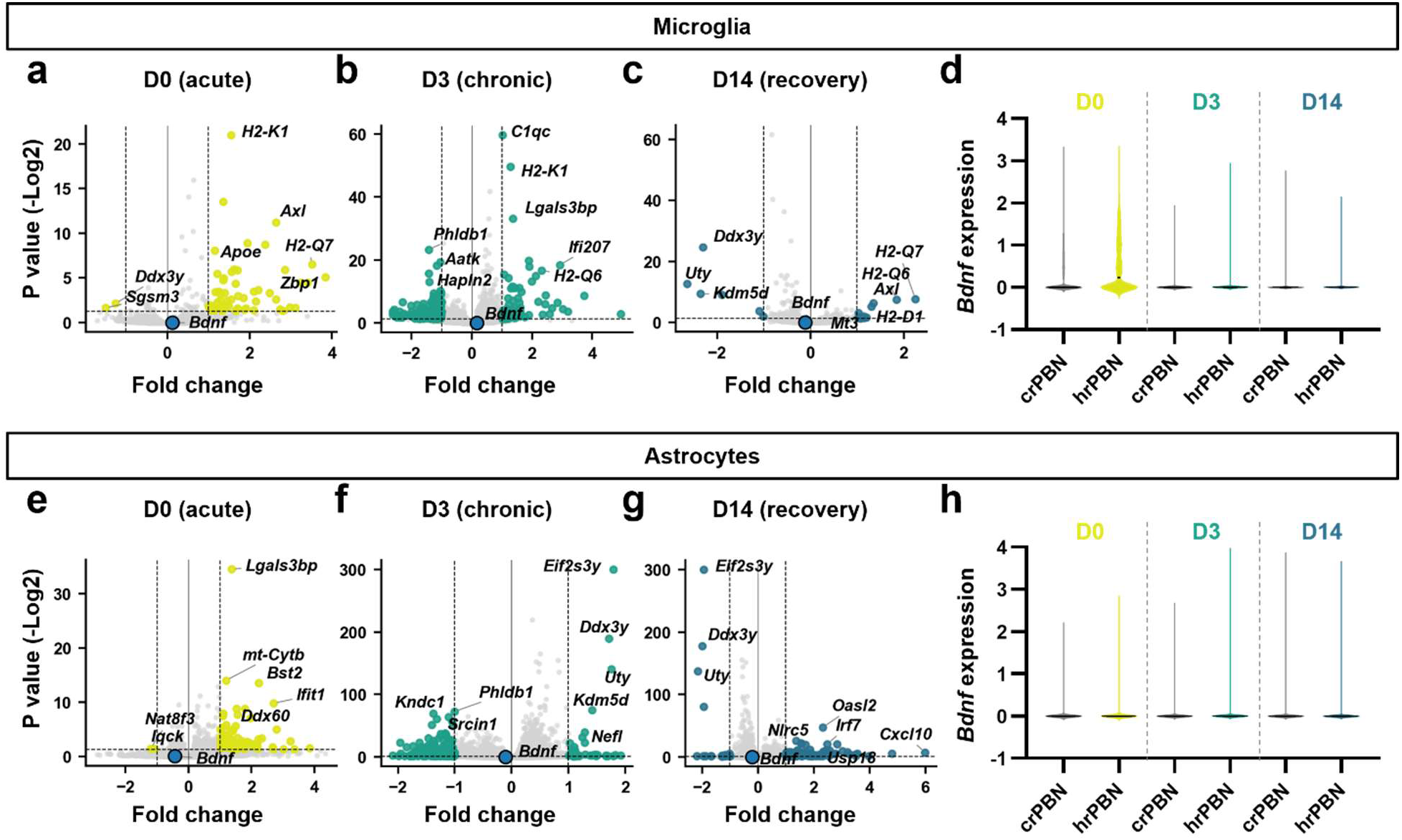
DEGs in microglia and astrocytes. **a-c,** Volcano plots of DEGs between crPBN and hrPBN in microglia at (a) D0, (b) D3, and (c) D14. **d,** Violin plots of *Bdnf* expression in microglia across phases. Black lines indicate median. **e-g,** Volcano plots of DEGs between crPBN and hrPBN in astrocytes at (e) D0, (f) D3, and (g) D14. **h,** Violin plots of *Bdnf* expression in astrocytes across phases. Black lines indicate median. In **d** and **h**, *Bdnf* expression between crPBN and hrPBN was compared by DESeq2 on pseudobulk profiles (Wald test) with Benjamini-Hochberg-adjusted P values. Note that the comparisons for microglia *Bdnf* could not be performed because of the low fraction of *Bdnf*-expressing cells.

**Extended Data Fig. 8.**
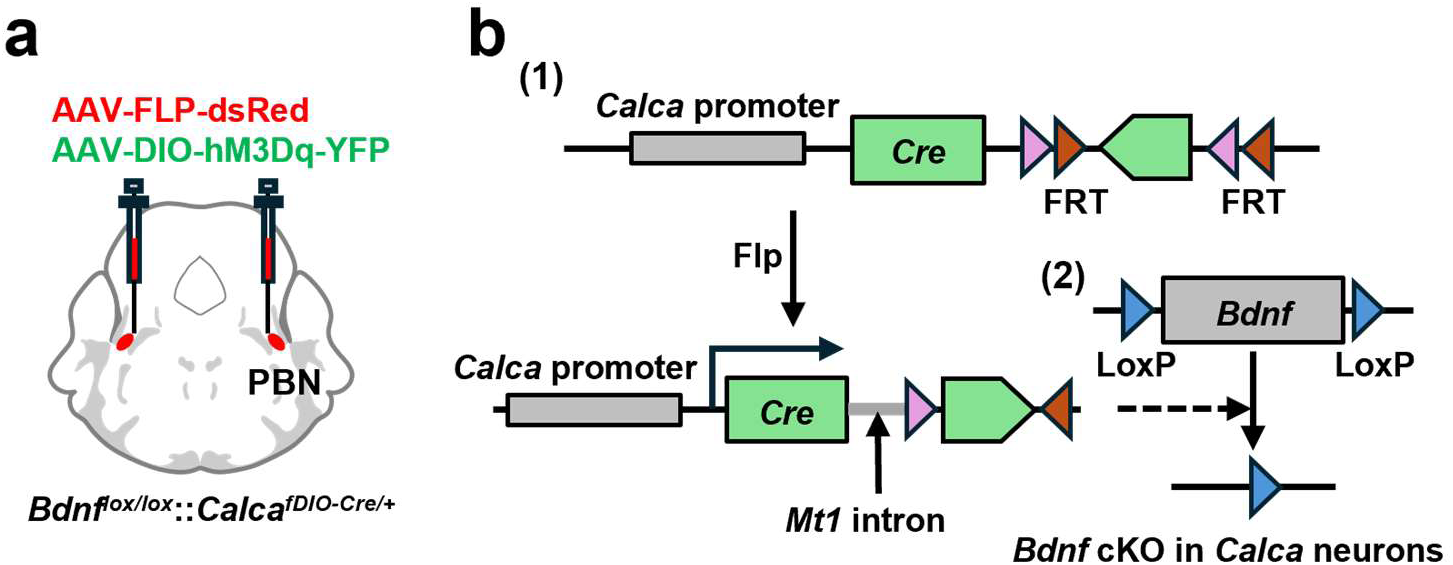
Schematic illustration of conditional knockout of *Bdnf* in *Calca* neurons. **a,** Conditional *Bdnf* knockout strategy. **b,** *Bdnf* knockout process. (1) Action of AAV-FLP:dsRed restores expression of 2-exon Cre recombinase driven by *Calca* promotor. (2) Induced Cre recombinase deletes *Bdnf* gene.

**Extended Data Fig. 9.**
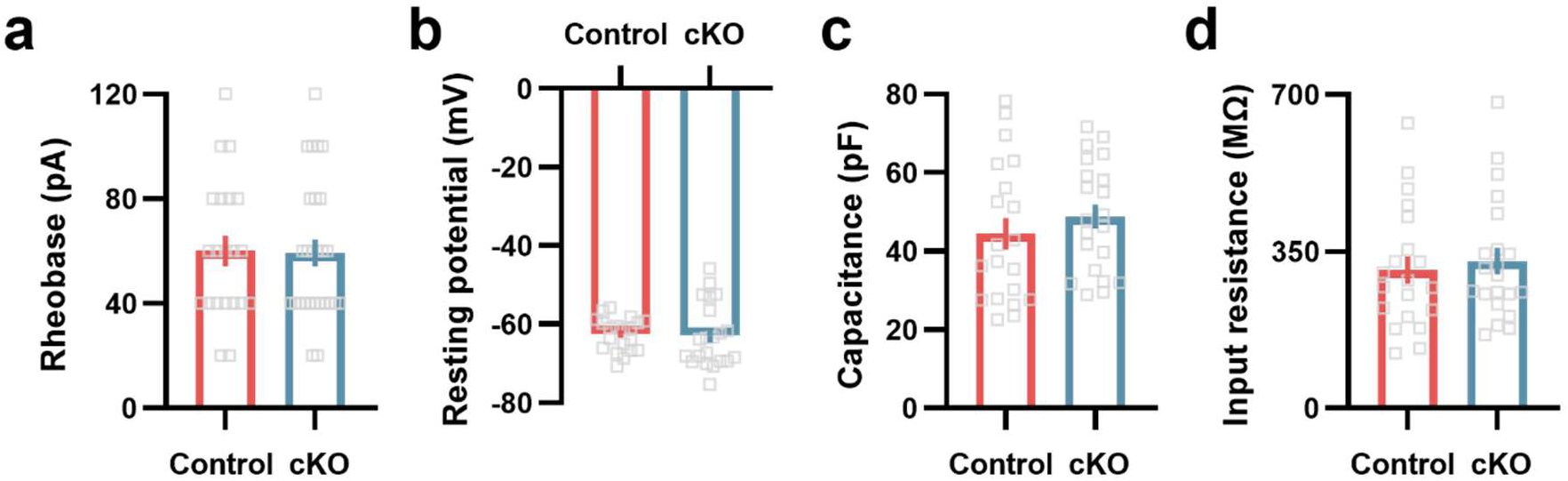
Membrane intrinsic properties of *Bdnf* control and cKO *Calca* neurons. **a,** Rheobase (n = 21 for *Bdnf* control and n = 27 for *Bdnf* cKO *Calca* neurons, Unpaired t-test). **b,** Resting potential. **c,** Capacitance. **d,** Input resistance. **b-d,** n = 20 for *Bdnf* control and n = 21 for *Bdnf* cKO *Calca* neurons. Unpaired t-test.

**Extended Data Figure 10.**
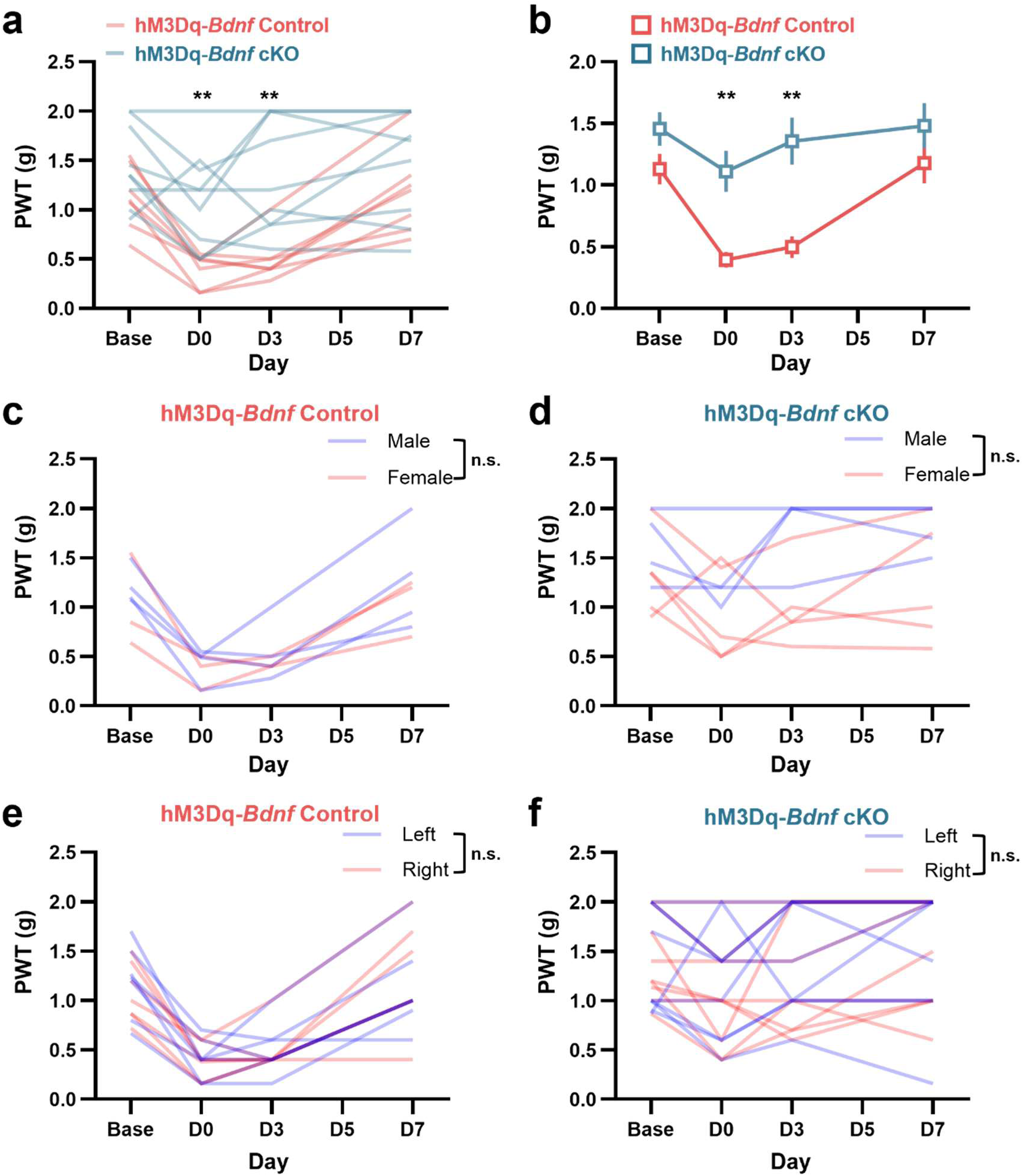
Behavioral responses of *Bdnf* control and cKO mice in *Calca* activation-driven persistent allodynia. **a-b,** Paw-withdrawal threshold of (a) individual animals and (b) average measured by von Frey test in *Calca* neuron-driven allodynia (n = 7 for control and n = 9 for cKO mice). c-d, Paw-withdrawal threshold in between male and female of *Bdnf* control and cKO groups (n = 4 for male and n = 3 for female in control; n = 4 for male and n = 5 for female in cKO group). e-f, Paw-withdrawal threshold in between left and right paw of *Bdnf* control and cKO groups (n = 7 for control and n = 9 for cKO mice). All data between the groups were compared by Two-way RM ANOVA. Data are means ±SEM.

**Extended Data Fig. 11.**
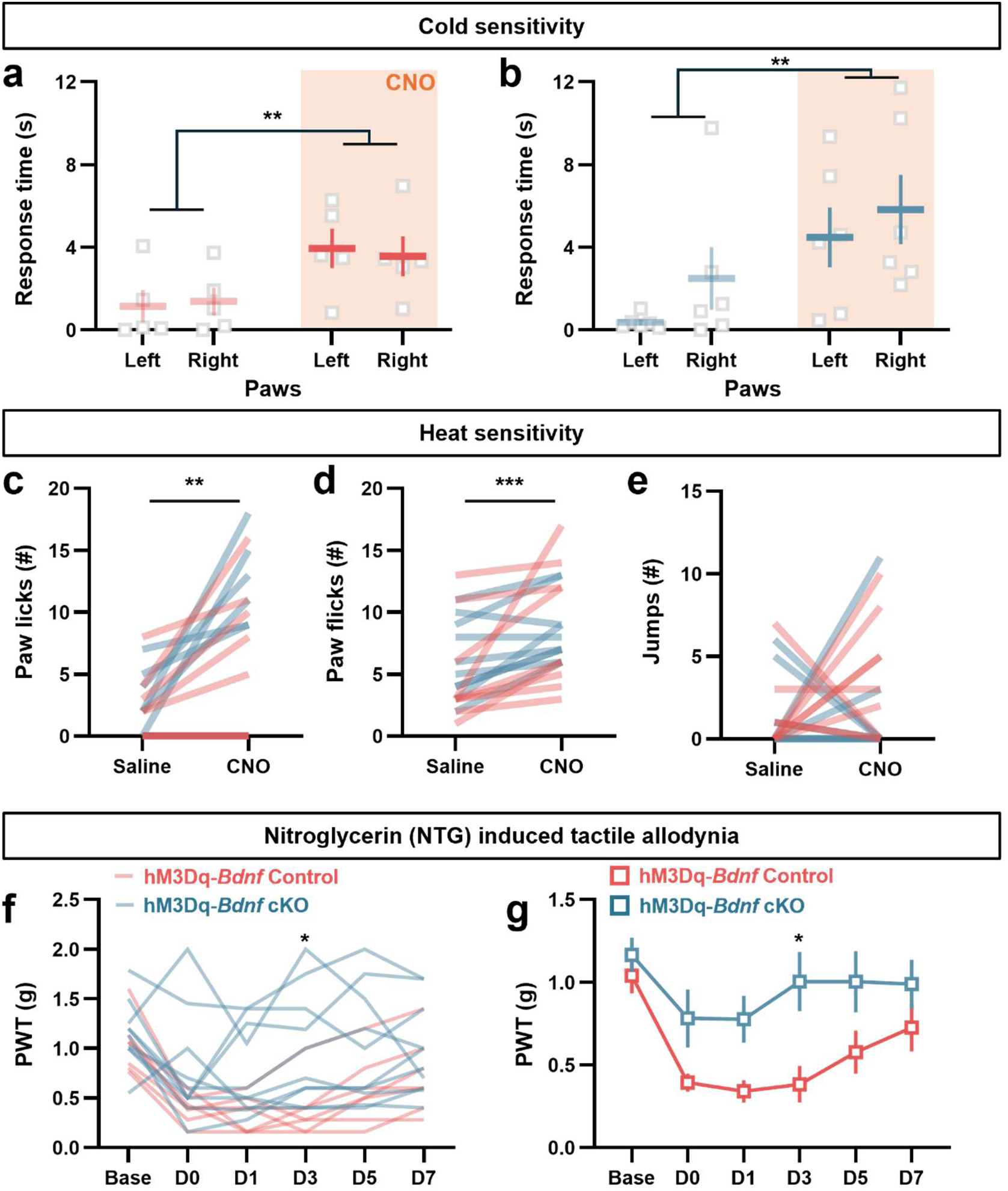
Behavioral responses of *Bdnf* control and cKO mice in temperature sensitivity test and NTG-induced migraine model. **a-b,** Cold sensitivity measured by acetone assay of each paw across individual control (a) and cKO (b) mice (n = 5 for control and n = 6 for cKO mice, Unpaired t test). **c-e,** Behavioral responses of individual animals during the hot plate test such as number of paw licks (c), paw flicks (d) and jumps (e). n = 9 for control and n = 10 for cKO mice (Unpaired t test). **f-g**, Paw-withdrawal threshold across individual *Bdnf* control and cKO mice and average response in NTG-induced migraine like pain (n = 7 for control and n = 10 for cKO mice, Two-way RM ANOVA).

## SUPPLEMENTARY TABLES

**Supplementary Table 1.** *Calca* neurons across neuronal clusters. The table shows the percentage of *Calca* expressing cells within each cluster. *Tac1* and *Nts* are known to be co-expressed with *Calca*, whereas *Foxp2* and *Tacr1* show little or no overlap with *Calca*. Clusters are ordered according to the percentage of *Calca* expressing cells.

| Cluster | # cells | % *Calca* | % *Tac1* | % *Nts* | % *Foxp2* | % *Tacr1* |
| --- | --- | --- | --- | --- | --- | --- |
| 14 | 1236 | 31.39 | 61.3 | 31.6 | 2.6 | 25.6 |
| 1 | 8669 | 31.17 | 61.2 | 29.7 | 1.7 | 1.7 |
| 11 | 2510 | 6.49 | 37.2 | 10.1 | 15.9 | 27.6 |
| 4 | 5864 | 5.51 | 67.8 | 18.0 | 1.7 | 1.9 |
| 16 | 497 | 3.42 | 23.3 | 7.8 | 11.5 | 12.3 |
| 2 | 8385 | 1.32 | 18.2 | 6.6 | 2.3 | 18.3 |
| 13 | 1629 | 1.23 | 7.6 | 2.8 | 3.1 | 8.1 |
| 6 | 5165 | 0.91 | 11.8 | 2.3 | 1.8 | 1.4 |
| 8 | 4648 | 0.60 | 28.5 | 4.5 | 1.4 | 9.3 |
| 7 | 4835 | 0.54 | 53.5 | 2.4 | 8.7 | 13.7 |
| 12 | 2443 | 0.53 | 14.6 | 16.2 | 66.5 | 13.7 |
| 0 | 9573 | 0.27 | 14.8 | 3.3 | 14.6 | 5.2 |
| 3 | 7351 | 0.14 | 5.8 | 2.6 | 15.1 | 46.9 |
| 10 | 4208 | 0.14 | 12.9 | 3.7 | 30.3 | 27.9 |
| 9 | 4306 | 0.12 | 8.2 | 4.0 | 2.6 | 11.5 |
| 5 | 5642 | 0.11 | 9.6 | 3.3 | 68.1 | 11.5 |
| 15 | 837 | 0.00 | 17.4 | 4.8 | 8.5 | 18.5 |

**Supplementary Table 2.**
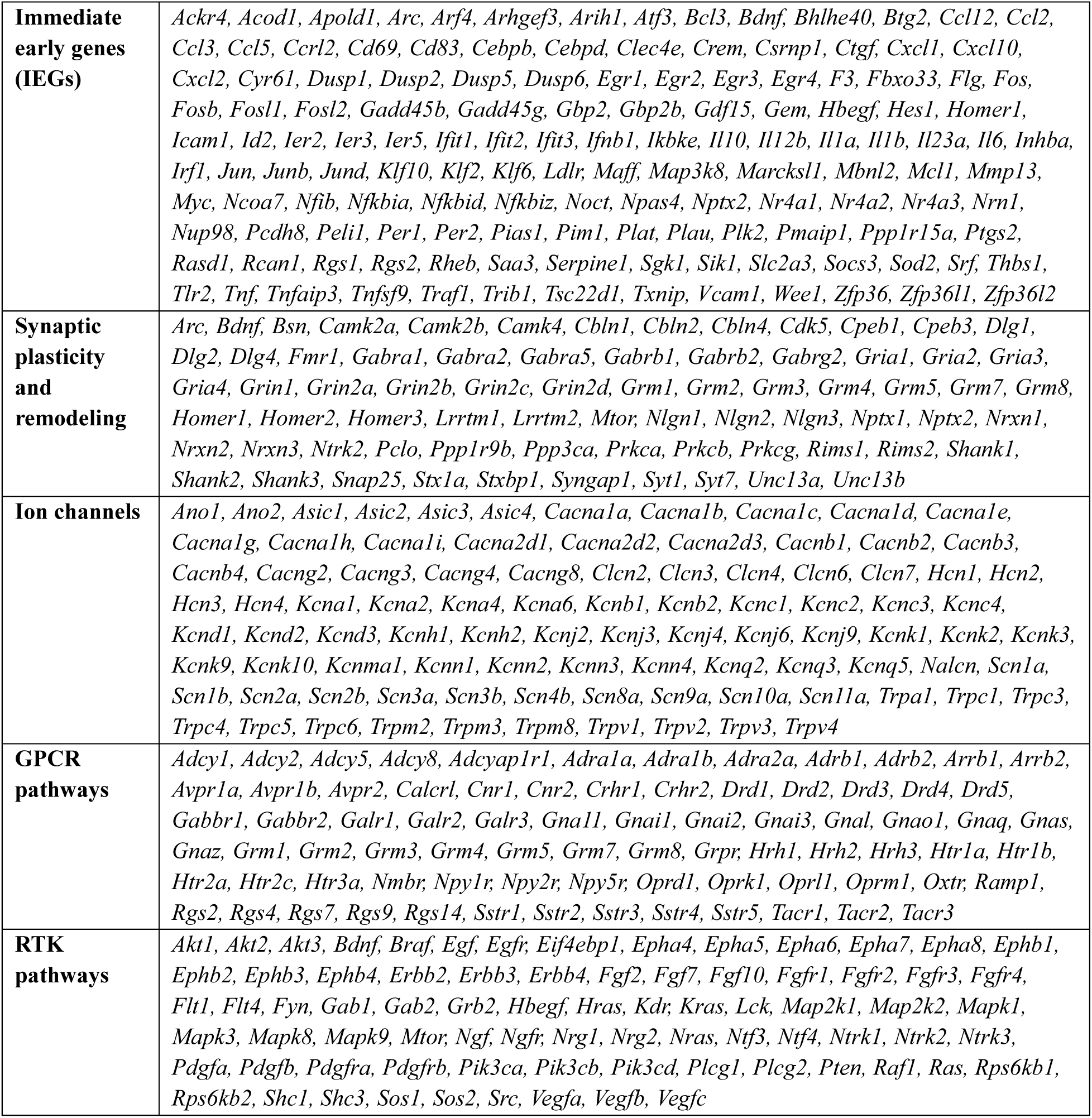
Lists of genes are used in DEG analysis.

## Notes

### Competing Interest Statement

The authors have declared no competing interest.

### Summary of Updates

This revision corrects figure legend errors in Figures 1g and 3b, clarifies gene labeling in Figure 3e, fixes in-text citations to Extended Data Fig. 11 and Supplementary Table 2, and makes minor text corrections throughout.

